# Differential Mast Cell Outcomes Are Sensitive to FcεRI-Syk Binding Kinetics

**DOI:** 10.1101/147595

**Authors:** Samantha L. Schwartz, Cédric Cleyrat, Mark Olah, Peter Relich, Genevieve Phillips, William S. Hlavacek, Keith A. Lidke, Bridget S. Wilson, Diane S. Lidke

## Abstract

Crosslinking of IgE-bound FcεRI triggers multiple cellular responses, including degranulation and cytokine production. Signaling is dependent on recruitment of Syk via docking of its dual SH2 domains to phosphorylated tyrosines within the FcεRI immunoreceptor tyrosine-based activation motifs. Using single molecule imaging in live cells, we directly visualized and quantified the binding of individual mNeonGreen-tagged Syk molecules as they associated with the plasma membrane after FcεRI activation. We found that Syk colocalizes transiently to FcεRI and that Syk-FcεRI binding dynamics are independent of receptor aggregate size. Substitution of glutamic acid for tyrosine between the Syk SH2 domains (SykY130E) led to an increased Syk-FcεRI off-rate, loss of site-specific Syk autophosphorylation, and impaired downstream signaling. CRISPR-Cas9 engineered cells expressing only SykY130E were deficient in antigen-stimulated calcium release, degranulation and production of some cytokines (TNF-a, IL-3) but not others (MCP-1, IL-4). We propose that kinetic discrimination along the FcεRI signaling pathway occurs at the level of Syk-FcεRI interactions, with key outcomes dependent upon sufficiently long-lived Syk binding events.

**Summary:** Schwartz et al. use single molecule imaging to quantify the transient nature of FcεRI-Syk interactions in live mast cells. A functional mutation that increases Syk off-rate leads to loss of site-specific Syk phosphorylation and impaired signaling, highlighting the importance of finely tuned protein interactions in directing cellular outcomes.

## Introduction

The family of multi-chain immunorecognition receptors (MIRRs), including the high affinity IgE receptor (FcεRI), the B cell receptor (BCR) and the T cell receptor (TCR), trigger a wide array of signaling outcomes critical for immune cell function including cell survival, release of inflammatory mediators and cytokine production. A distinguishing feature of the MIRRs is their lack of intrinsic kinase activity, rendering them reliant on the recruitment and activation of non-receptor tyrosine kinases for signaling (Sigalov, 2005). For FcεRI and BCR, antigen engagement results in phosphorylation of accessory chain Immunoreceptor Tyrosine-based Activation Motifs (ITAMs) (Johnson et al., 1995) by the Src family kinases Lyn and Fyn, followed by the recruitment and activation of the tyrosine kinase Syk. The parallel signaling cascade of the TCR relies on sequential engagement of the Src family kinase, Lck, and the Syk-related kinase Zap70.

As the only two members of a kinase subfamily, Syk and Zap70 share structural similarities that regulate kinase activity through conformational state changes. Key features include tandem SH2 domains that are joined via a linker (Interdomain A; I-A), and a kinase domain (KD) connected via a second linker region (Interdomain B; I-B) (Au-Yeung et al., 2009). Interactions between the linker domains and the kinase domain create an auto-inhibited conformation (Deindl et al., 2007); this closed state is likely the predominant conformation of Syk/Zap70 in a resting cell. Transitions to an open conformation facilitate binding of the tandem SH2 domains to dually-phosphorylated ITAMs and free the kinase domain for activity (Johnson et al., 1995). The ITAM-docked open conformation also exposes a number of autophosphorylation sites, as well as tyrosines that are substrates for Src kinases (Arias-Palomo et al., 2009; Chen et al., 2011; Geahlen, 2009; Sada et al., 2001). Phosphorylation of Syk at these sites promotes kinase activation (Tsang et al., 2008) and provides docking sites for distinct downstream signaling molecules (Simon et al., 2005). Phosphotyrosines are also implicated in structural modifications of Syk. For example, phosphorylation of I-B tyrosine residues in both Syk and Zap70 favors the open state (Brdicka et al., 2005) and recent studies have shown that phosphorylation of these residues in Zap70 is associated with longer dwell times on the TCR (Klammt et al., 2015). Phosphorylation of key residues in the catalytic domain of Syk also enhances enzymatic activity (Carsetti et al., 2009). It is important to note that, despite their similarity, these two closely related kinases also have important distinctions. Notably, Zap70 activation by the TCR is reliant on the Src family kinase Lck, as well as the protein phosphatase CD45, whereas Syk can reconstitute TCR signaling without these two players (Chu et al., 1996). Reliable and quantitative measures that capture receptor-kinase interaction dynamics are needed to understand the full range of mechanisms underlying the similarities and differences between these essential immune kinases (Palacios and Weiss, 2007; Turner et al., 2000).

This work focuses on the dynamics of Syk recruitment to FcεRI following aggregation of the receptor by multi-valent antigen. Immuno-electron microscopy studies have shown that FcεRI crosslinking leads to a dramatic increase in the amount of Syk associated with the receptor at the plasma membrane (Wilson et al., 2000). However, FcεRI aggregates can vary in size and mobility as a function of antigen dose or valency (Andrews et al., 2009; Mahajan et al., 2014). Therefore, establishing the relationship between receptor aggregate size and signaling efficiency is of keen interest (Wilson et al., 2011). Mathematical modeling studies have predicted that membrane receptor clustering can lead to enhanced signaling output by promoting multiple rebinding events (Das et al., 2009; Radhakrishnan et al., 2012). On the other hand, membrane topographical features or post-translational modifications that reduce diffusion-mediated access or dwell time at the ITAM domain could negatively influence signaling. Thus, while structural and biochemical studies have provided information on the sequence of events needed for Syk activation in this signaling cascade, many questions remain concerning the timing and extent of Syk recruitment needed to efficiently propagate signaling. We performed Single Particle Tracking (SPT) using Total Internal Reflection Fluorescence (TIRF) microscopy to quantify the dynamics of Syk recruitment to the membranes of gene-edited RBL-2H3 cells, whose endogenous Syk expression was ablated and reconstituted with either wild type (WT) or mutant Syk expressed as a chimeric mNeonGreen (mNG) fusion protein. Immunological and biochemical assays were used to determine the impact of changes in FcεRI-Syk dynamics on mast cell functional responses.

We find that Syk binding at the mast cell membrane is best described as a mixture of lifetimes characterized by a fast off-rate (*k_f_* = 2.6 s^−1^) and slow off-rate (*k_s_* = 0.62 s^−1^), indicating a population of both short-lived and long-lived binding events. Aggregation of FcεRI leads to a marked increase in the fraction of trajectories characterized by *k_s_* compared to *k_f_*, and treatment with the Lyn specific inhibitor Dasatinib drastically reduces this *k_s_* fraction. These results indicate that *k_s_* characterizes specific recruitment of Syk to phosphorylated FcεRI. Based upon two-color imaging, Syk-FcεRI colocalization is sustained through rapid exchange with the pool of cytosolic Syk. The importance of the longer-lived interactions in signal propagation is shown by introduction of a Y130E mutation within the I-A domain of Syk. Phosphorylation of Y130 is proposed as a form of negative feedback regulation since it has been shown to destabilize binding of Syk tandem SH2 domains to phosphorylated ITAMs (pITAM) (Feng and Post, 2015; Zhang et al., 2008). We find that Syk-Y130E is still recruited to FcεRI aggregates, but its interactions are more transient (*k_s_* = 0.87 s^−1^) and markedly less efficient at transphosphorylation. In cells expressing only the Syk-Y130E mutant form of Syk, mast cell degranulation and specific cytokine production (TNFα, IL3) is impaired but remarkably, production of MCP-1 and IL4 is retained.

In previous work it has been shown that the kinetics of ligand-receptor binding impact signaling events and cellular responses (Liu et al., 2001; McKeithan, 1995; Suzuki et al., 2014; Torigoe et al., 2007). FcεRI signal transduction therefore, has classically been considered to be controlled at the step of receptor aggregation according to the principles of kinetic proofreading (Hopfield, 1974). Here, we find that propagation of a subset of cellular responses is similarly sensitive to the off-rate of Syk recruited to aggregated receptors. These data highlight the importance of finely tuned protein-protein interactions in directing cellular outcomes.

## Results

### Syk recruitment to FcεRI is transient

We began by generating a Syk knockout sub-line of RBL-2H3 cells (Syk-KO) through CRISPR-Cas 9-mediated editing. These cells were then reconstituted with WT Syk expressed as an mNG fusion protein (Syk^mNG^). Syk-mediated downstream signaling in these cells was fully rescued, including restoration of degranulation responses downstream of FcεRI aggregation (Fig. S1). To follow Syk^mNG^ recruitment to activated FcεRI, we next primed cells with fluorescent anti-DNP IgE (Alexa Fluor 647-conjugated IgE; AF647-IgE) and then challenged cells with the multivalent antigen, DNP-BSA (0.1 μg/mL). Confocal images in Fig. 1A and Supplementary Video 1 show the expected increase in FcεRI clustering that accompanies 5 min of antigen-simulated aggregation and the formation of signaling patches (Wilson et al., 2000). Receptor crosslinking also led to the accumulation of Syk^mNG^ at the membrane, where it co-localized with FcεRI aggregates (Fig. 1A; Supplementary Video 1). The recruitment of Syk^mNG^ to the plasma membrane was lost in cells pretreated with Dasatanib, which selectively inhibits Src and Abl tyrosine kinases (Lombardo et al., 2004) (TIRF images, Fig. 1B). Further proof that the observed binding events are specific to FcεRI aggregation is shown in Fig. 1C, where the disruption of FcεRI aggregates by monovalent DNP-lysine also results in rapid dissociation of Syk^mNG^ clusters. After the addition of DNP-lysine, single molecules could be resolved near the original cluster prior to release into the cytosol (Fig. 1C, bottom). The time course for this series of events can be viewed in Video 2. The apparent persistence of Syk at the membrane over this short period may reflect interactions with phosphorylated FcεRI monomers or other membrane components (possibly LAT or other substrates) that persist for a short time after the dissolution of FcεRI aggregates.

**Figure 1.**
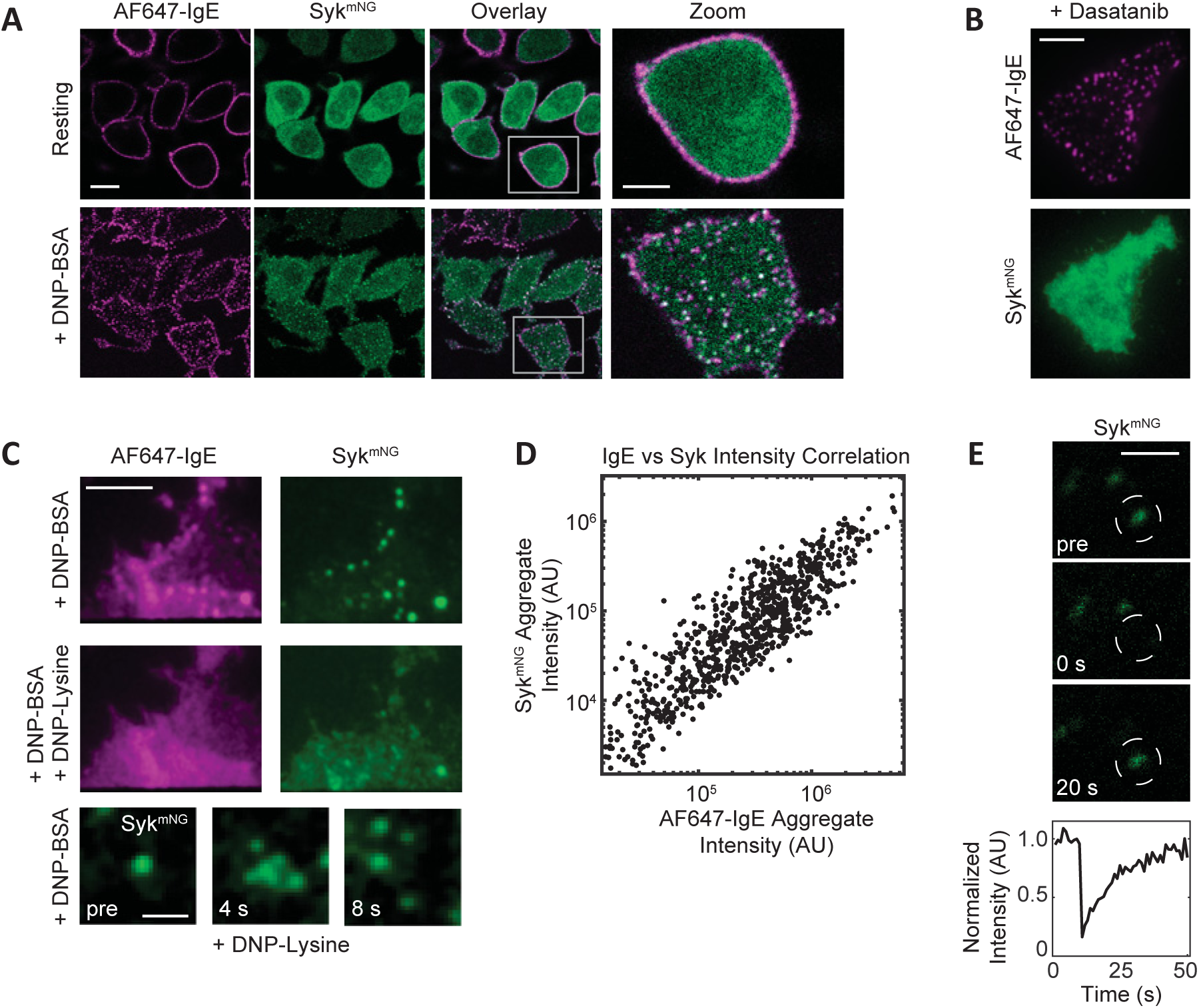
Syk Recruitment to FcεRI. Syk-KO RBL cells expressing Syk^mNG^ (green) were primed with AF647-IgE (magenta) and imaged after crosslinking with 0.1 μg/mL DNP-BSA. (A) Sample images from a confocal time series showing the redistribution of AF647-IgE-FcεRI and Syk^mNG^ upon FcεRI stimulation (see also Video 1). Resting cross-section shows homogeneous distribution of AF647-IgE-FcεRI at the plasma membrane and Syk^mNG^ in the cytosol. Upon crosslinking (5 min), FcεRI aggregation and Syk^mNG^ colocalization is readily seen at the adherent cell surface. Scale bar, 10.3 µm. White boxes in the ‘Overlay’ panels are enlarged in the ‘Zoom’ panels. Scale bar, 2 µm. (B) Treatment with 1 µM Dasatinib results in a loss of Syk^mNG^ recruitment (bottom) to FcεRI aggregates (top). Images of the adherent cell membrane acquired in TIRF. Scale bar, 5 µm. (C) Both FcεRI and Syk^mNG^ aggregates (top) are disrupted upon addition of 100 µM monovalent DNP-lysine (middle). Scale bar, 5 µm. Selected images from a time series (bottom) show the dispersion of an individual Syk aggregate within seconds of DNP-lysine addition. Individual Syk^mNG^ molecules can be seen diffusing away from the original diffraction-limited aggregate (see also Video 2). Images acquired in TIRF. Scale bar, 1 gm. (D) Plot of positive correlation between Syk^mNG^ and AF647-IgE intensity within each AF647-IgE aggregate. (E) Selected images from a confocal time series before and after photobleaching (at t=0 s) of an individual Syk^mNG^ aggregate. Scale bar, 1 µm. Bottom curve quantifies the rapid recovery of mNG fluorescence intensity within the bleached region (white circles).

We next examined the recruitment capacity of FcεRI aggregates by comparing receptor aggregate size and density with Syk^mNG^ accumulation. Using two-color TIRF imaging, AF647-IgE images were first segmented by creating an intensity mask to identify individual receptor aggregates, from which, corresponding AF647-IgE and Syk^mNG^ intensities were determined. The linear correlation of the IgE-FcεRI and Syk^mNG^ intensities per aggregate seen in Fig. 1D indicates that as receptor aggregates increase in size, more Syk^mNG^ is recruited. Finally, we assessed the dynamics of FcεRI-Syk interactions using fluorescence recovery after photobleaching (FRAP). Syk^mNG^ co-localized with FcεRI aggregates demonstrated rapid fluorescence recovery within 20 s (Fig. 1E) while the FcεRI does not (data not shown). These results reveal that the observed Syk^mNG^ aggregation is not stable in time, but actually an accumulation of many transient binding events.

### Direct measurements of Syk binding dynamics

To directly measure the off-rate of Syk binding, we applied single molecule imaging to visualize thousands of Syk^mNG^ binding events in living cells. Using TIRF microscopy, we were able to observe and track single Syk^mNG^ molecules as they associated with the adherent surface of the plasma membrane (Video 3). We selected our imaging frame rate (100 ms exposure time) to minimize the contribution of fast moving Syk^mNG^ molecules in the cytosol and selectively capture those Syk^mNG^ proteins that reduce mobility when bound to the membrane (Fig. 2A, left). In this scenario, the track length of individual Syk^mNG^ proteins reflects the binding lifetime (Fig. 2A, right). As shown in the cumulative probability plots in Fig. 2B, we found that the distribution of track lengths shifted to longer duration after FcεRI activation. To extract the underlying Syk off-rates (*k*), we fit the distribution of track lengths assuming an exponential binding process and compared fits across different models (see Methods for details). We found the simplest model consistent with the data was a two-component model with a fast off-rate (*k_f_*) of ~2.6 s^−1^ and a slower off-rate (*k_s_*) of ~0.62 s^−1^ (Fig. 2C). We interpret the faster off-rate to represent non-productive binding events (i.e., sampling of monophosphorylated ITAMs or other membrane partners) and the slow off-rate to characterize the specific Syk docking events that are correlated with FcεRI signaling. Consistent with this interpretation, we found that the *k_s_* fraction markedly increased upon FcεRI activation, from 5% to 23% (Fig. 2D). The increase in the *k_s_* fraction with stimulation is blocked when cells are pretreated with the Src family kinase inhibitor Dasatanib (Fig. 2D). These results indicate that the longer-lived events are specifically associated with Syk binding to the FcεRI pITAM. Of note, we found that there was a significant population (5%) of *k_s_* Syk^mNG^ binding events in resting cells, which was essentially ablated (<2%) with Dasatinib treatment. This suggests that some basal phosphorylation of FcεRI occurs in the absence of crosslinking to recruit Syk, although we cannot fully exclude the possibility of specific Syk binding to other Src (or Abl) substrates at the membrane.

**Figure 2.**
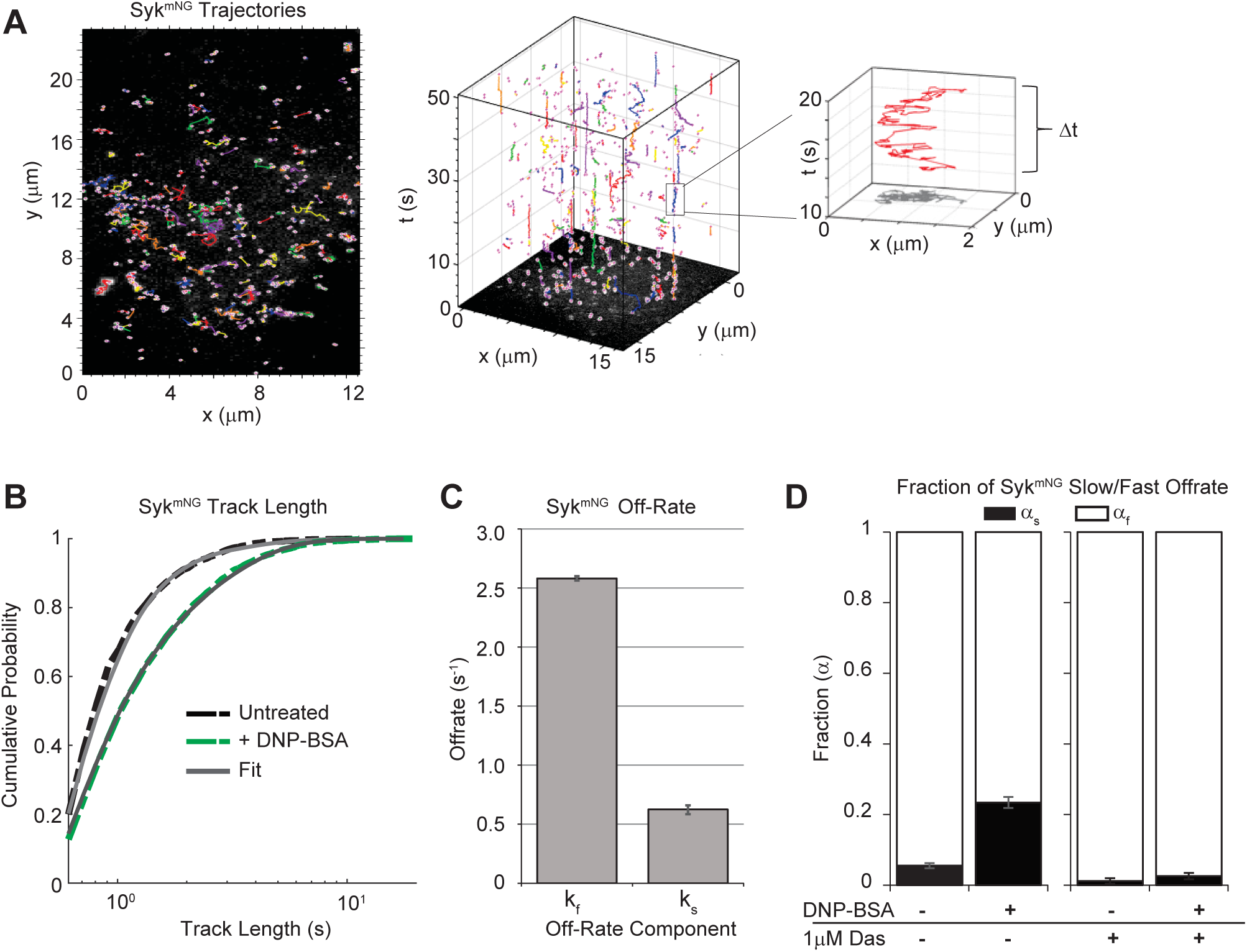
Quantifying FcεRI-Syk interactions using single particle tracking of Syk^mNG^. Single Syk^mNG^ molecules were tracked using TIRF microscopy as they bound and dissociated from the membrane. (A) Example of Syk^mNG^ trajectories detected at the plasma membrane (left). Projection of trajectories in the time dimension (middle) shows the trajectory lengths. Enlargement of a single trajectory that lasts ~10 s (right). (B) Cumulative probability distribution of Syk^mNG^ trajectory lengths before (black) and 4-5 min after addition of 1 μg/mL DNP-BSA (green). Solid grey lines represent the fit to the data. (C) Bar graph depicts fast and slow off-rates found when fitting the distributions in (B). See Methods for details. (D) Fraction of the slow off-rate component α_s_ increases from ~5% to 23% after addition of 1 μg/mL DNP-BSA. Treatment with 1 µM Dasatinib (Das) reduces α in both resting and activated cells. Error bars in (C-D) are a 68% credible interval as described in Methods. See also Table 1.

**Table 1.**
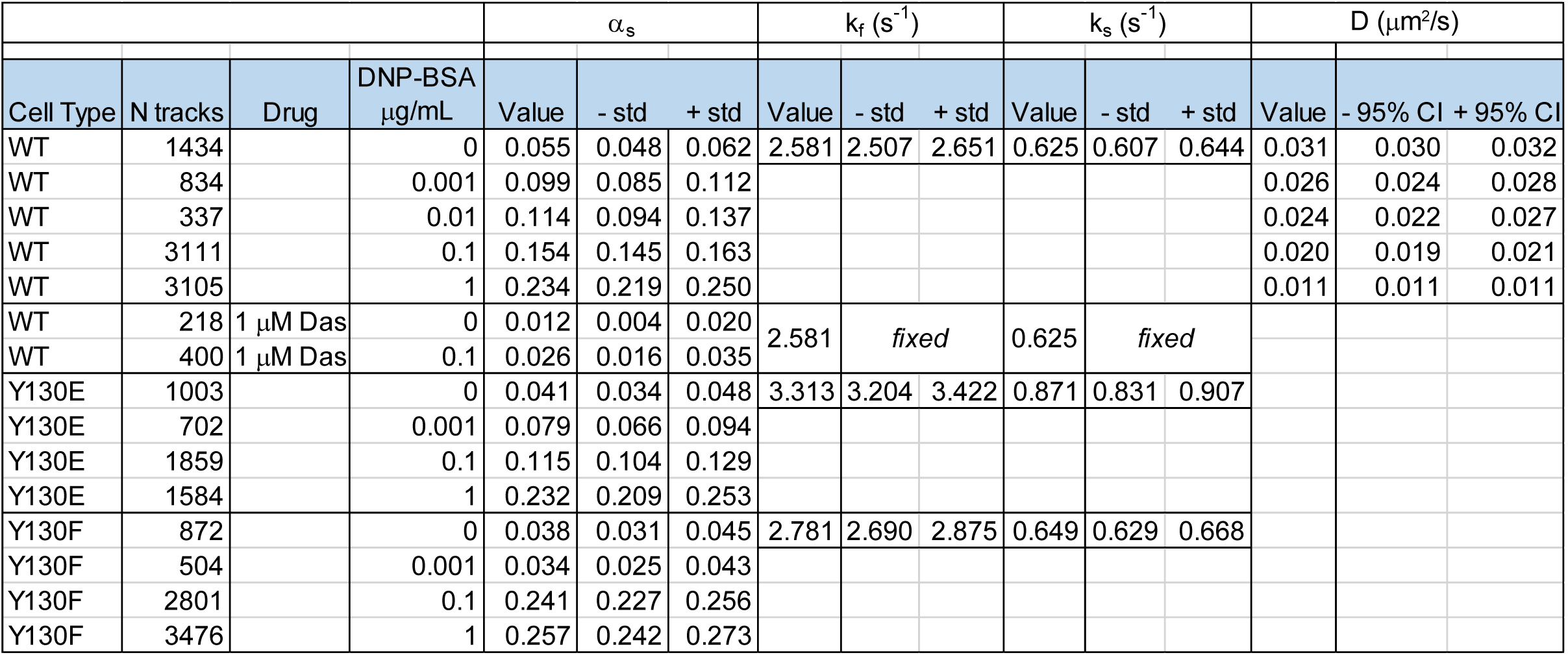
Measured Syk off-rates and diffusion coefficients. Off-rate parameter values found from fitting single molecule track lengths to a two-component geometric distribution mixture model, with both off-rate parameter values globally constrained across cell type in Syk-KO cells expressing Syk^mNG^-WT (WT), Syk^mNG^-Y130E (Y130E), or Syk^mNG^-Y130F (Y130F). Data for each condition displayed with cell type, the number of trajectories used in the analysis, if addition of 1 µM Dasatinib treatment, and dose of DNP-BSA (μg/mL). The estimated fast off-rate (k_f_), the slow off-rate (k_s_), and the relative fraction of slow off-rate (α) are reported (Value) along with a standard error below (-std) or above (+std) based on a 68% credible interval (see Methods). For the WT +1 μM Das conditions, the off-rate values were *fixed* to the WT untreated rate. The estimated diffusion coefficients (D) for the Syk^mNG^-WT are reported in the right most column along with the lower (-95%CI) and upper (+95% CI) 95% confidence intervals.

### FcεRI-Syk interaction dynamics are independent of antigen dose or aggregate size

The extent and time course of FcεRI aggregation is a feature of antigen dose and valency (Andrews et al., 2009). To establish the aggregation profile of FcεRI after crosslinking with DNP-BSA, we used dSTORM (van de Linde et al., 2011) super-resolution imaging. Imaging was performed in TIRF, which specifically evaluates antigen-mediated aggregation at the adherent surface. Results in Fig. 3A,B report clustering of AF647-IgE bound receptors after 5 min of treatment over a range of DNP-BSA doses (0.001 −1 μg/ml). The comparison in Fig. 3A, between the diffraction-limited images at left and the dSTORM images at right, illustrates that the ~10 nm localization accuracy of this super-resolution method enables the visualization and quantification (Fig. 3B) of aggregation across the full dose response range. Aggregation is observed even at the lowest doses (0.001 and 0.01 μg/mL) where changes in the diffraction-limited image are not readily discernable.

**Figure 3.**
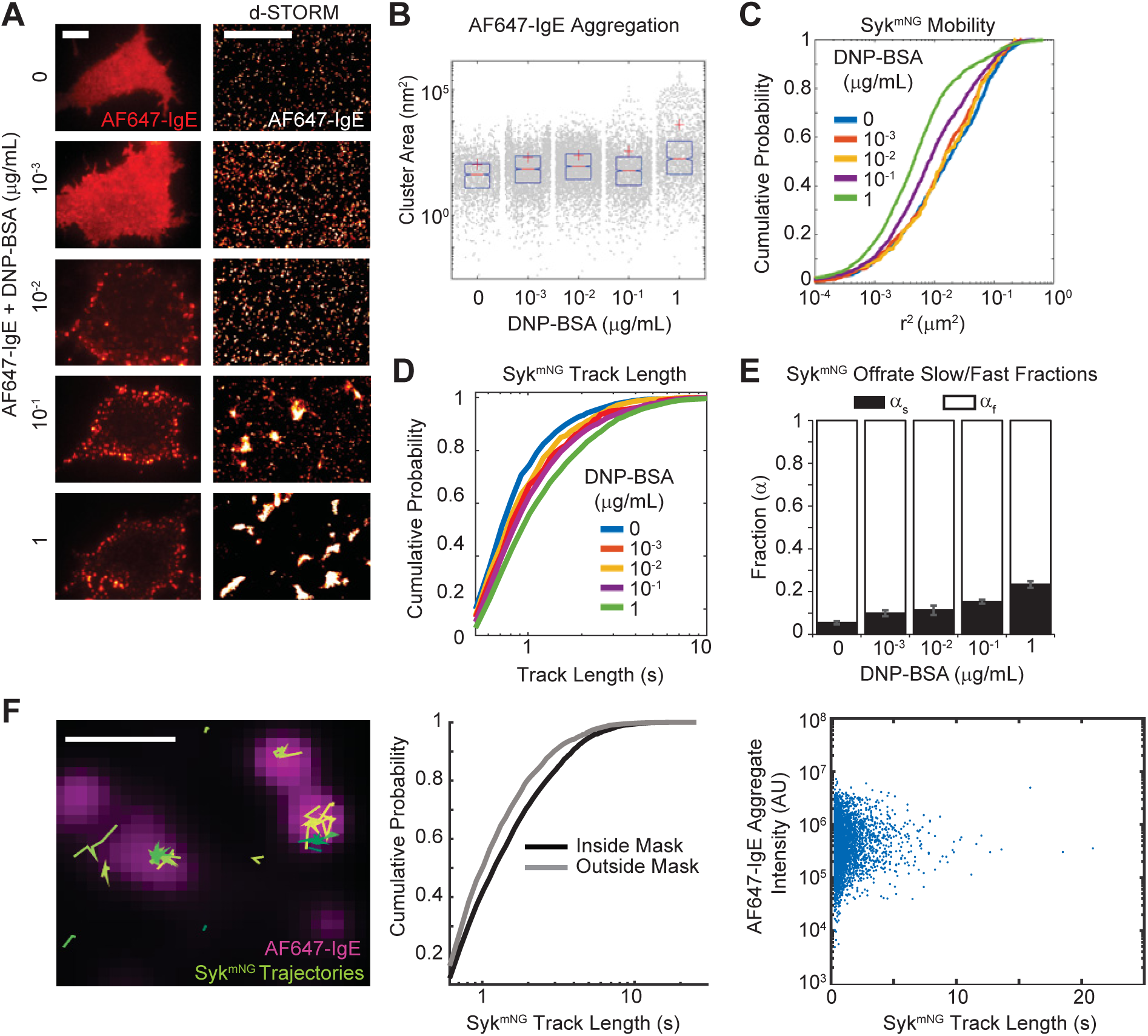
FcεRI-Syk off-rate is independent of antigen dose or aggregate size. (A) AF647-IgE labeled RBL cells crosslinked for 5 min with indicated DNP-BSA concentration and imaged (left) at the adherent surface in TIRF (Scale bar, 5 µm) and using (right) dSTORM (Scale bar, 2 µm). (B) Clustering of localizations in dSTORM images from (A) using a Hierarchical clustering algorithm (Matlab, The MathWorks, Inc.). Cluster sizes are shown as grey dots and the distribution is summarized by the mean (red cross), the median (red line) and the 25th and 75th percentiles (blue box). Kolmogorov-Smirnov tests show significant (p<0.01) differences between resting (0 ng) and other DNP-BSA doses. (C) Mobility of Syk^mNG^ represented as the cumulative distribution of squared displacements (r^2^, ΔT = 0.3 sec or 3 frames) for each DNP-BSA dose. See Table 1 for values. (D) Cumulative distribution of Syk^mNG^ track lengths for each DNP-BSA dose. (E) The fraction of the slow off-rate component ((α_s_) increases with DNP-BSA dose. Data collected between 1-5 min after the addition of DNP-BSA. Error bars are a 68% credible interval as described in Methods. See also Table 1. (F) Comparison of Syk^mNG^ trajectory localization with AF647-IgE aggregates in cells stimulated with 0.1 μg/mL DNP-BSA for up to 5 min. Left: Overlay of Syk^mNG^ trajectories (green lines) with AF647-IgE aggregates (magenta). Scale bar, 1 µm. See also Video 4. Middle: Comparison of Syk^mNG^ track length distributions inside and outside the AF647-IgE mask. A Kolmogorov-Smirnov test shows the distributions are significantly different (p<0.01). Right: No correlation is observed between Syk^mNG^ track length and the intensity of the AF647-IgE aggregate it co-localizes to.

We showed previously that FcεRI aggregates become essentially immobile at the highest antigen doses (≥ 0.1 μg/ml) (Andrews et al., 2009). Since both small (mobile) and large (immobile) complexes were shown to be signaling competent (Andrews et al., 2009; Shelby et al., 2014), it was important to determine if there were differences in Syk recruitment across all crosslinking conditions. Fig. 3C and Table 1 show that the mobility of Syk^mNG^ recruited to the membrane is reduced as antigen dose increases, following the same trend as the antigen-induced changes in FcεRI mobility (Andrews, 2009). Fig. 3D shows that the distribution of track lengths shifts toward longer duration as a function of antigen dose. Fitting these distributions across the dose response again supported a model in which the rate constants remain unchanged, rather it is the fraction of long-lived tracks (α_s_) that increase with the concentration of antigen (Fig. 3E, Table 1). These data suggested that the interaction lifetime of Syk at the membrane is independent of the mobility of receptor aggregates or their cluster size.

To confirm this conclusion, we used two-color imaging to simultaneously correlate receptor aggregate size with Syk^mNG^ dwell times (Fig. 3F; Video 4). We chose an intermediate dose of DNP-BSA concentration (0.1 μg/mL), where a range of AF647-IgE-FcεRI aggregate sizes can be seen on the cell surface (Fig. 3A,B). For each Syk trajectory, we compared its track length with the total intensity of the corresponding AF647-IgE-FcεRI aggregate it co-localized to (Fig. 3F). Comparison of Syk^mNG^ dwell times inside and outside the aggregates showed that most long-lived events are associated with FcεRI aggregates (Fig. 3F, middle). No correlation between aggregate intensity (i.e., size) or Syk^mNG^ dwell time was found (Fig. 3F, right). Thus, we interpret these results to indicate that signaling is associated with an increase in the number of long-lived Syk binding events rather than changes in binding lifetime.

### Phosphomimetic mutation at Y130 increases the Syk off-rate

Considering the observation that FcεRI-Syk interaction kinetics are consistent across all DNP-BSA crosslinking conditions, we next sought to experimentally alter the lifetime of Syk on FcεRI pITAMs by introducing mutations in tyrosine 130 in the Syk I-A domain. Located between the two tandem SH2 domains, Y130 phosphorylation reportedly lowers the affinity of Syk for pITAMs (Isaacson, 1997; Zhang et al., 2008). We reconstituted Syk-KO cells with mNG-tagged versions of two previously described Syk mutants (Isaacson, 1997) (Fig. S2A): 1) Y130E, a phosphomimetic mutation (Syk^mNG^-Y130E); and 2) Y130F, which cannot become phosphorylated (Syk^mNG^-Y130F). These mutants were also introduced into parental RBL-2H3 cells (Fig. S2A). As shown in Fig. 4A, each of the reconstituted Syk-KO cell lines showed robust membrane accumulation of Syk^mNG^ after stimulation (0.1 μg/mL DNP-BSA) that colocalized with AF647-IgE-FcεRI aggregates. Quantification of these images, shown in Fig. 4B, was performed in the same manner as Fig. 1D. The Y130E and Y130F mutants of Syk^mNG^ were recruited to FcεPRI to a similar extent as wild type Syk^mNG^. Thus, despite poor recovery of the Y130E mutant in co-immunoprecipitation studies with the BCR (Isaacson, 1997; Zhang et al., 2008), association of Syk^mNG^-Y130E with FcεRI aggregates is not grossly impaired.

**Figure 4.**
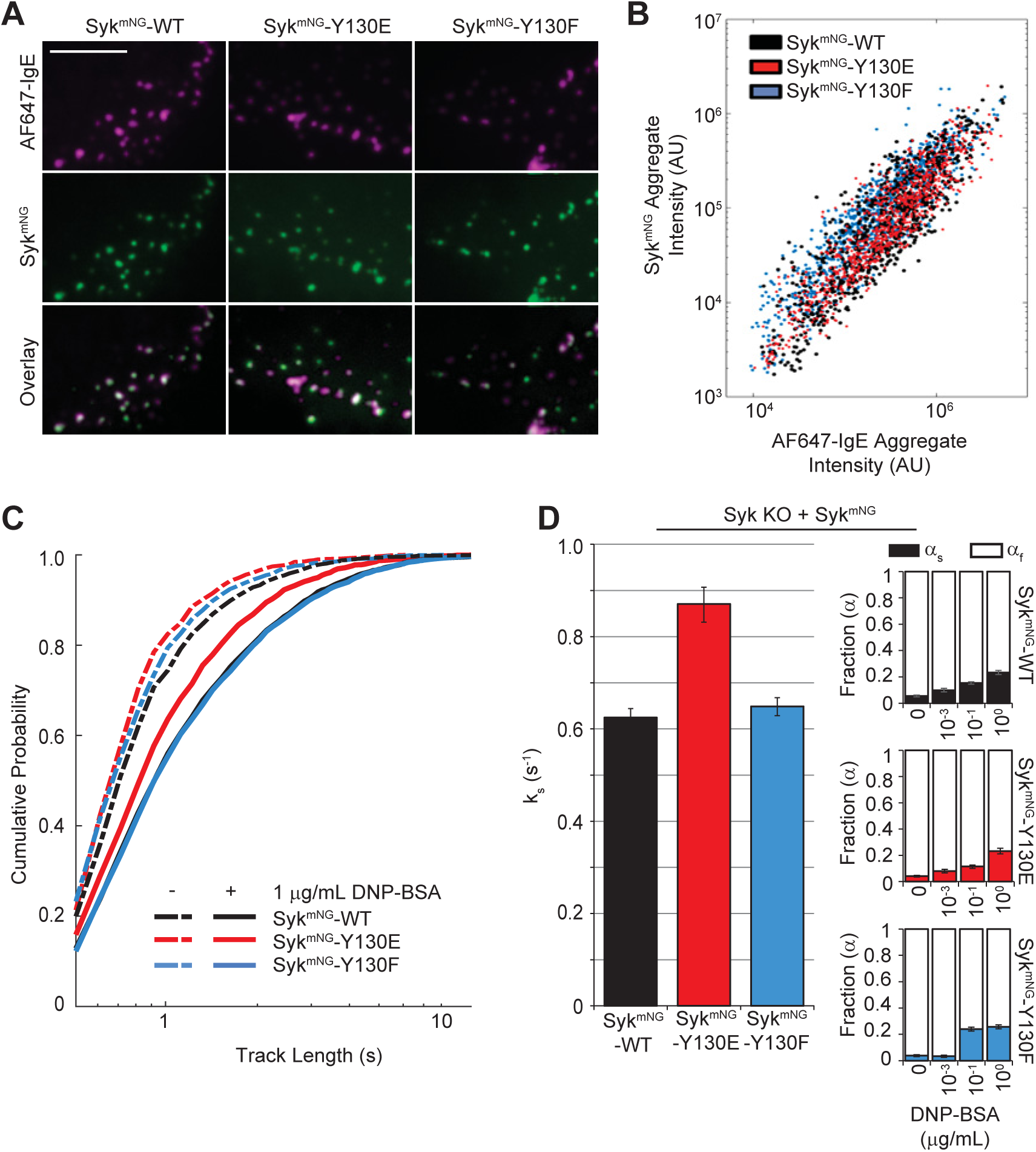
Syk^mNG^-Y130E exhibits a faster FcεRI off-rate. (A) TIRF images of AF647-IgE (magenta) and Syk^mNG^ (green) membrane localization in Syk-KO cells reconstituted with Syk^mNG^-WT (left), Syk^mNG^-Y130E (middle) or Syk^mNG^-Y130F (right) after 4-5 min stimulation with 0.1 μg/mL DNP-BSA. Scale bar, 5 µm. (B) Quantification of FcεRI recruitment capacity for Syk. Individual aggregates of AF647-IgE were masked and the total intensity within the mask for both the AF647-IgE and Syk^mNG^ channels is plotted per aggregate. Recruitment is similar for Syk^mNG^-WT and each mutant. (C) Cumulative probability distributions of trajectory lengths for Syk^mNG^-WT, -Y130E and -Y130F both before (dashed lines) and after (solid lines) addition of 1 μg/mL DNP-BSA. (D) Bar graph depicts the slow off-rate (k_s_) found when fitting the distributions in (C). Fraction of slow off-rate component (α_s_) increases with DNP-BSA dose for Syk^mNG^-WT and each mutant (right). Error bars are a 68% credible interval (see Methods).

We next considered whether these mutations could alter the interaction dynamics of Syk at the membrane. Using the Syk^mNG^-Y130E and Syk^mNG^-Y130F stable cells lines, we used TIRF microscopy to image single Syk^mNG^ binding events for both mutant forms of Syk. In the resting state, both the Y130E and Y130F mutants have a similar track length distribution to that of WT (Syk^mNG^-WT) (Fig. 4C, dashed lines). Interesting differences were observed however, upon activation with DNP-BSA. Although Syk^mNG^ -WT and Syk^mNG^-Y130F showed the expected increase in track length distribution with FcεRI activation, track lengths of Syk^mNG^-Y130E did not shift to the same extent. These distributions were best fit to a two-component model with a modified off-rate, *ks*, for Y130E mutants of 0.87 s^−1^, compared to 0.62 s^−1^ for Syk^mNG^-WT and 0.65 s^−1^ for Syk^mNG^-Y130F (Fig. 4D, Table 1). An increased off-rate for Syk^mNG^-Y130E was also observed in the parental cells (Fig. S3A,B). The fraction of *k_s_* for all three Syk isoforms exhibited a similar increase as a function of DNP-BSA dose (Fig. 4D insets), consistent with the idea that higher antigen doses are associated with an increase in the number of long-lived Syk binding events.

### Mast cell responses are sensitive to Syk lifetime on FcεRI

Mast cells are known to release histamine and other granule contents within minutes of FcεRI stimulation, followed by transcriptional upregulation of cytokine and chemokine production (Metcalfe et al., 1997). To explore whether the subtle dynamic changes we observed for the Syk-Y130E off-rate have functional consequences, we evaluated the impact on FcεRI-mediated release of inflammatory mediators. Results in Fig. 5A and Fig. S1 show that Syk-KO cells reconstituted with either Syk^mNG^-WT or Syk^mNG^-Y130F have comparable degranulation responses to antigen stimulation (0.1 μg/ml DNP-BSA, 30 min; black, blue bars). In contrast, cells reconstituted with Syk^mNG^-Y130E were incapable of secreting preformed mediators under the same stimulating conditions (red bar, Fig. 5A). To ensure that Syk^mNG^-Y130E reconstituted cells were competent for degranulation, replicate samples were also treated with Ionomycin (1 μM) plus Phorbol 12-myristate 13-acetate (PMA). Results are reported in Fig. S2B showing that Syk-Y130E cells do release granule contents in response to stimuli that bypass the receptor to elevate calcium levels. Additionally, we found that kinase activity *in vitro* was similar for all three Syk variants (Fig. S2C); thus, impaired degranulation in the Syk^mNG^-Y130E expressing cells cannot be explained by altered kinase activity.

**Figure 5.**
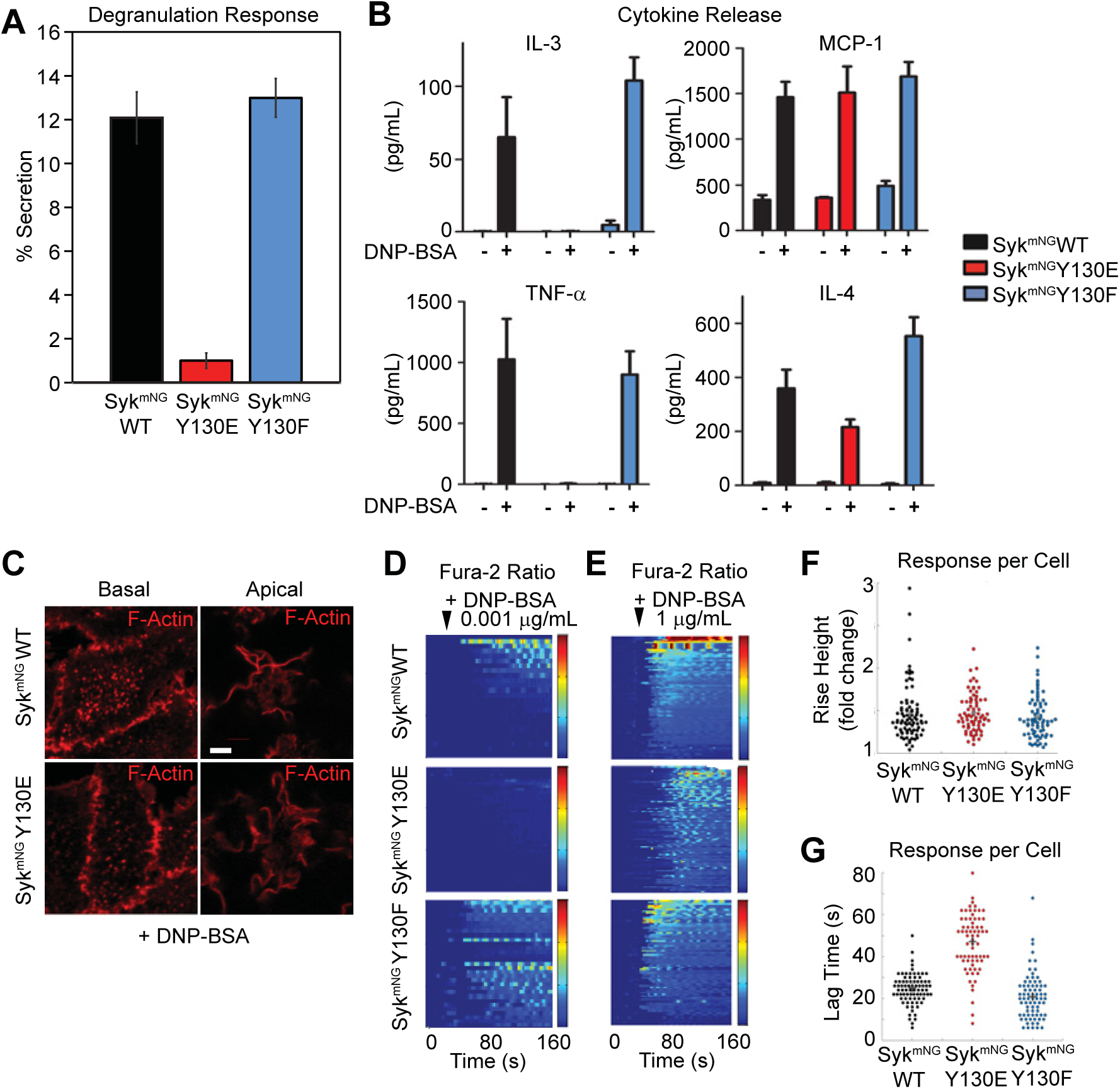
Key mast cell outcomes are impaired in Syk^mNG^-Y130E cells. (A) Degranulation mea-sured by relative β-hexosaminidase released after 30 min of incubation with 0.1 μg/mL DNP-BSA in Syk-KO cells reconstituted with Syk^mNG^-WT, Syk^mNG^-Y130E or Syk^mNG^-Y130F. (B) Comparison of cytokine concentration in cell media of Syk-KO cells reconstituted with Syk^mNG^-WT, Syk^mNG^-Y130E, or Syk^mNG^-Y130F before and after 3 hrs stimulation with 0.1 μg/mL DNP-BSA. Results were repeated at least three times for all four cytokines. Bar plots represent mean and standard deviation of technical replicates in one representative sample preparation. (C) The formation of actin plaques at the basolateral surface (left) and ruffling on the apical surface (right) in response to 0.1 μg/mL DNP-BSA treatment in both Syk^mNG^-WT and Syk^mNG^-Y130E reconstituted Syk-KO cells. Filamentous actin labeled with phalloidin-AF647 (red). Scale bar, 2 µm. (D-E) Heat maps of relative changes in intracellular calcium concentration upon addition of either a (D) low dose (0.001 μg/mL) or (E) high dose (1 μg/mL) of DNP-BSA. Each row represents the ratio of Fura-2 emission using 350 nm/380 nm excitation for a single cell over time. Ratio color bar scale range, 1-2 fold increase: Blue-Red. (F) Relative increase in Fura-2 ratio per cell after addition. (G) Time between antigen addition and response for each cell. (F-G) Stimulated with 1 μg/mL of DNP-BSA and calculated as described in Methods.

We also evaluated release of newly formed mediators from each of the three Syk-reconstituted cell lines during 3 hrs of simulation with 0.1 μg/ml DNP-BSA. Results are shown in Fig. 5B for the cytokines IL-3, TNFα and IL-4, as well as the chemokine MCP-1 (CCL2). In Syk^mNG^-WT and Syk^mNG^-Y130F reconstituted cells, antigen stimulation led to production of all four factors (Fig. 5B). Syk^mNG^-Y130E cells, on the other hand, completely lost the ability to produce TNFα and IL-3 while production of MCP-1 was normal, or somewhat attenuated, compared to WT or Y130F expressing cells. IL-4 production in Syk^mNG^-Y130E cells was variable, but detectible in 2 of 4 samples.

FcεRI activation is also known to induce PI3K-dependent ventral cell ruffling and dorsal actin plaque formation (Barker et al., 1995; Pfeiffer and Oliver, 1994). Fig. 5C shows that these responses are comparable in Syk-KO cells reconstituted with either Syk^mNG^-WT or the Syk^mNG^-Y130E mutant. Since PI3K-mediated production of PtdIns(3,4,5)P3 allosterically enhances PLCγ activity in this system (Barker et al., 1999; Smith et al., 2001), we next performed ratiometric imaging of calcium signaling in Fura-2 loaded single cells. Heat maps in Fig. 5D,E report the composite profiles of calcium responses in at least 30 cells per condition following the addition of antigen. Stimulation with low antigen dose (0.001 μg/ml DNP_24_-BSA) induced measurable calcium responses in Syk^mNG^-WT (33 out of 71 cells) or Syk^mNG^–Y130F (37 out of 72 cells) reconstituted cells, consistent with previous observations. By comparison, Syk^mNG^-Y130E expressing cells were markedly impaired, with only ~10% of cells (8 out of 84 cells) demonstrating any measurable calcium response to low antigen dose. Notably, Syk^mNG^-Y30E expressing cells were capable of initiating a calcium response after challenge with high antigen dose (1 μg/ml), as shown in Fig. 5E. However, while the amplitude of the calcium response was similar between the three cell types at high antigen dose (Fig. 5F), the onset of calcium flux was delayed by ~20 s in the Syk-Y130E expressing cells (Fig. 5G).

### Syk Phosphorylation Kinetics are finely tuned to ITAM interaction times

Results shown above indicate that Syk-Y130E strikingly fails to support a number of key mast cell outcomes (degranulation and TNFα and IL-3 release), whereas other responses downstream of FcεRI activation were unaffected (MCP-1 production, ruffling). These data suggest that the reduced interaction times observed for Syk-Y130E correlate with differences in phosphorylation linked to specific arms of the FcεRI signaling network. Fig. 6A shows representative Western blotting results comparing the extent of phosphorylation at four critical tyrosine residues (Y317, Y342, Y346, Y519/520) within wild type and mutant Syk^mNG^ after 5 min stimulation with 0.1 μg/mL DNP-BSA. Results from multiple experiments are represented in the bar graphs of Fig. 6A. These data show that both Syk^mNG^-WT and Syk^mNG^-Y130F were strongly phosphorylated at all four tyrosine sites in response to antigen, while in cells reconstituted with Syk^mNG^-Y130E phosphorylation at these tyrosines was varied. Two of the sites in the I-B linker region (Y317, Y346) are known substrates of Lyn (Keshvara et al., 1998; Sanderson et al., 2010), although they can also be trans-phosphorylated by adjacent Syk molecules (Tsang et al., 2008). The other two sites (Y519/520 and Y342) are critical Syk autophosphorylation sites. In cells reconstituted with Syk^mNG^-Y130E, phosphorylation at Y317 and Y346 was unimpaired while phosphorylation of Y519/520 and Y342 was markedly inhibited: we observed a two-fold reduction in phosphorylation at Y519/520 and a five-fold reduction in phosphorylation at Y342 (Fig. 6A, right). These results were confirmed in parental RBL-2H3 cells expressing these constructs (Fig. S3C), demonstrating that the presence of endogenous wild type Syk cannot rescue Syk^mNG^-Y130E phosphorylation at these two sites.

**Figure 6.**
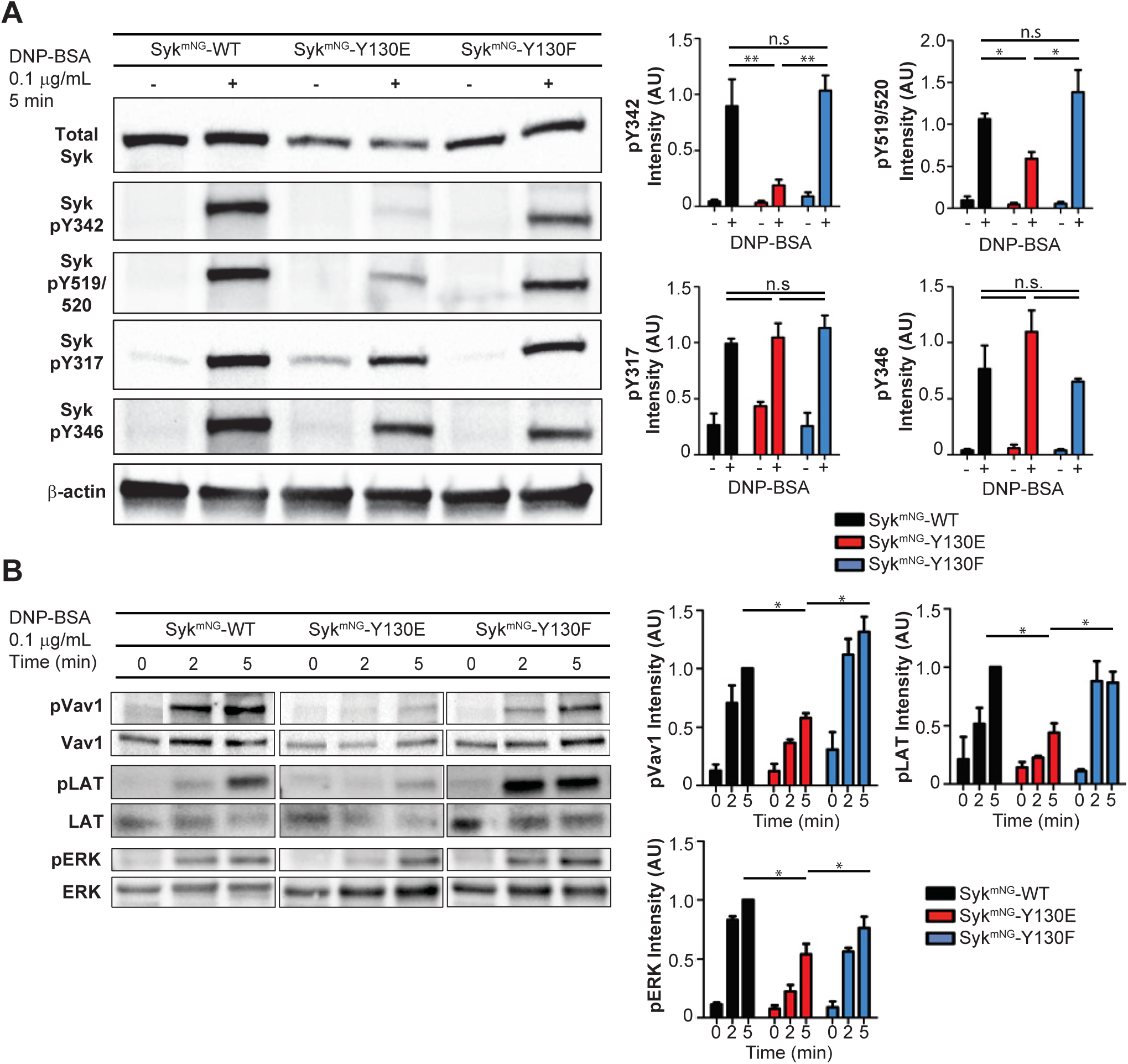
Phosphorylation kinetics of Syk and downstream signaling partners. (A) Western blot detection of the Syk phosphorylation profile in Syk-KO cells reconstituted with Syk^mNG^-WT, Syk^mNG^-Y130E, or Syk^mNG^-Y130F in response to stimulation with 0.1 μg/mL DNP-BSA for 5 min. Bar plots (right) represent mean and standard deviation of relative Syk phosphorylation level from at least three experiments. Phosphorylation of tyrosines associated with Syk auto-catalytic activity (pY519/520 and pY342) is significantly reduced (ttest, *p<0.05,**p<0.01) in Syk^mNG^-Y130E. No significant difference (n.s) in phosphorylation is seen at sites susceptible to Lyn catalytic activity (pY317, pY346). (B) Western blot detection of the phosphorylation time course of downstream signaling molecules Vav1, LAT and ERK. Cells were stimulated with 0.1 μg/mL DNP-BSA for 0, 2, or 5 min. Bar plots (right) represent mean and standard deviation of the relative phosphorylation levels from three experiments (ttest, *p<0.05).

Decreased phosphorylation at specific Syk autophosphorylation sites is predicted to impair docking of Syk binding partners and/or inhibit their Syk-mediated activation by phosphorylation. Western blotting results in Fig. 6B report representative changes in tyrosine phosphorylation profiles for two Syk substrates downstream of FcεRI activation: LAT (Wilson et al., 2004) and Vav1 (Margolis et al., 1992). We also evaluated phosphorylation of Erk as a read out of the Ras-MAPK pathway (Graham et al., 1998). In cells expressing Syk^mNG^-WT, LAT phosphorylation peaked at 5 min after stimulation with 0.1 μg/ml DNP24-BSA. The onset of LAT phosphorylation was consistently faster in cells reconstituted with Syk^mNG^-Y130F (n=3), peaking at 2 min rather than 5 min. Thus, this mutant is equal to (or slightly better) than Syk^mNG^-WT at coupling with LAT. In contrast, LAT phosphorylation was delayed in Syk^mNG^-Y130E and reached only 50% of Syk^mNG^-WT levels by 5 min. We also found that antigen-stimulated Syk^mNG^-Y130E cells had a ~50% reduction in Vav1 phosphorylation compared with Syk^mNG^-WT cells, consistent with prior work identifying Syk pY342 as a key docking site for Vav1 (Deckert et al., 1996). Finally, phosphorylation of Erk under these conditions was also reduced in Syk^mNG^-Y130E expressing cells compared to cells reconstituted with either Syk^mNG^-WT or Syk^mNG^-Y130F. Taken together, these results illustrate that the reduced residency time for Syk^mNG^-Y130E translates to impaired autophosphorylation and impaired coupling to at least three downstream signaling branches.

## Discussion

Signaling networks are composed of a series of reactions bifurcating from ligand-bound receptors. The importance of key elements, including cellular colocalization of critical players and detailed biochemical reaction schemes, has laid the foundation for our modern understanding of signal transduction. Although these reactions are often depicted as a linear chain of events, the dynamic interplay of signaling partners and the rapid turnover of post-translational modifications and/or second messengers are critical components necessary to create a highly sensitive system. Studies of the high affinity IgE receptor (FcεRI) have contributed significantly to our understanding of the spatiotemporal aspects of cell signaling, due in part to the unique architecture of the signaling complexes formed after antigen-mediated crosslinking (Menon et al., 1986). At the onset of signaling, encounters between IgE-bound FcεRI monomers on membranes are diffusion-mediated (Andrews et al., 2009; Menon et al., 1986) and the resulting aggregates are influenced by the valency, affinity and dose of ligand. Since none of the subunits in the FcεRI αβγ_2_ tetramer bear intrinsic catalytic activity, propagation of signals is dependent on sequential recruitment of extrinsic kinases. The dually acylated Src family kinase, Lyn, is responsible for rapid ITAM phosphorylation (Eiseman and Bolen, 1992). Prior work has suggested that a fraction of Lyn is co-localized with FcεRI clusters in resting RBL-2H3 cells (Veatch et al., 2012; Wilson et al., 2000) and a small fraction may be pre-bound to the FcεRI β-subunit (Vonakis et al., 2001; Yamashita et al., 1994). ITAM phosphorylation supports recruitment of the dual SH2-containing Syk tyrosine kinase from the cytosol (Hutchcroft et al., 1992; Shiue et al., 1995).

With the knowledge that Syk translocation is essential for the release of inflammatory mediators from mast cells and basophils (Costello et al., 1996; Zhang et al., 1996), we monitored the dynamics of individual Syk molecules during recruitment to FcεRI in live cells. Recent studies have demonstrated the power of single molecule imaging methods in capturing the real-time interactions of membrane components with cytosolic binding partners, including clathrin assembly (Cocucci et al., 2012), the TCR’s recruitment of Zap70 (O’Donoghue et al., 2013), and Grb2 association with EGFR (Ichinose et al., 2006). These works have illustrated the inherent stochastic nature of binding events at the single molecule level, as well as the highly transient nature of protein interactions during active signaling and trafficking. Our SPT studies of Syk show that interactions with phosphorylated receptors are transient. We used the Syk trajectory lengths to extract a characteristic off-rate for the interaction of Syk at the membrane. From our fitting we found that the distribution of Syk trajectories was best characterized by a mixture of two rate constants, a non-specific, fast off-rate (*k_f_* = 2.6 s^−1^) and a slow off-rate (*k_s_* = 0.62 s^−1^) that was directly associated with FcεRI phosphorylation as it was ablated in Dasatinib-treated cells. The fraction of this slow off-rate component increased with higher DNP-BSA dose and IgE-FcεRI aggregation. Model comparison analysis of fits using a globally constrained *k_s_* value across all conditions versus a freely varying *k_s_* value revealed that our data was best fit to a model in which the *k_s_* value does not vary with antigen dose (only the fraction of trajectories characterized by *k_s_* varies with dose). This suggests that higher antigen doses and larger IgE-FcεRI aggregates do not alter the fundamental Syk-pITAM interaction kinetics, but increase the concentration of pITAMs available for Syk binding. Future work using different antigen geometries or affinities will be important to explore if other local receptor environments influence Syk-pITAM kinetics.

Mathematical models have led to the prediction that membrane receptor clustering can favor multiple rebinding events between receptors and their signaling partners (Das et al., 2009; Radhakrishnan et al., 2012). Given the limitations of frame rate at which our data was acquired, we considered the possibility that rapid rebinding events to receptors in the same complex aggregate might artificially inflate the observed Syk off-rate. However, we found no correlation between track length and size of receptor aggregates. One interpretation is that, in the unique case of Syk, rebinding may be prevented due to the sequence of conformational and post-translational changes.

ITAM-bearing receptors, such as the TCR, provided early motivation for use of kinetic proofreading models to study the dependence of signal transduction events on ligand-receptor binding kinetics (McKeithan 1995). In its simplest form, kinetic proofreading predicts that high affinity ligand-receptor bonds enable completion of a series of intermediate steps necessary to generate productive signals. This mechanism imposes constraints on signaling from non-specific interactions and provides sensitivity to differences in dose and affinity. However, the FcεRI system also provided early evidence that distal events, including cytosolic messengers, might provide an alternative means of kinetic discrimination for the generation of qualitatively different signals (Hlavacek et al., 2001; Liu et al., 2001). Prior work by others established conditions using low- and high-affinity ligands for crosslinking IgE-FcεRI complexes that achieved similar receptor phosphorylation levels but differential outcomes. For example, degranulation was reduced after challenge with low affinity ligands while chemokine production was unaffected or superior (Liu et al., 2001; Suzuki et al., 2014). Rivera and colleagues proposed that the signaling differences could be explained, at least in part, by differential recruitment of Src family kinases (Fgr), and the membrane scaffold, LAT2/NTAL (Suzuki et al., 2014). Both groups also reported a reduced phosphorylation of Syk with low affinity ligand stimulation. We show here that a single point mutation, Y130E, in Syk alters mast cell signaling in ways that are remarkably similar to stimulation with low affinity ligand (Liu et al., 2001; Suzuki et al., 2014). We used the same high affinity, high valency ligand across all our experiments comparing Syk^mNG^-WT and –Y130E cells. Therefore, we expect that FcεRI aggregation kinetics and geometry are consistent in our experiments and that recruitment of Src family kinase members is also unaltered. We found that the reduction in Syk-Y130E phosphorylation was site-specific. No change in the Lyn-dependent site Y346 (Sanderson et al., 2010) was observed, again consistent with similar levels of Lyn/Src kinase recruitment. However, sites subject to autophosphorylation (Sanderson et al., 2010) were variable: phosphorylation at Y317 was unchanged, Y519/520 was reduced two-fold and Y342 was reduced five-fold. This pattern suggests a hierarchy of phosphorylation events that is sensitive to FcεRI interaction conditions. It also explains, at least in part, the differential signaling outcomes. For example, pY342 has previously been shown to be critical for Syk to interact with and phosphorylate the downstream partner Vav1 (Deckert et al., 1996; Simon et al., 2005) while pY346 is a critical site for recruitment of Cbl ubiquitin ligase, implicated in negative regulation of FcεRI signaling (Keshvara et al., 1998; Lupher et al., 1998).

The correlation between Syk binding lifetime, phosphorylation kinetics and cellular outcome is consistent with the idea that the timing of interactions is critical to produce a fully modified protein. Previous *in vitro* experiments measured the binding of recombinant wild type and mutant Syk-SH2 domains to dually-phosphorylated TCR-ITAM peptides (Zhang et al., 2008). Based on these data, we expected as much as a 10-fold difference in binding lifetime for WT and Syk-Y130E. The difference that we measured in live cells is only 1.4 fold, which could be explained by conformational constraints of the intact Syk protein, spatial aspects of disulfide-linked full-length, dimeric γ-subunits and/or influence of the lipid bilayer on ITAM accessibility (Lopez et al., 2015). Because Syk also exhibits distinct affinities for particular ITAM sequences, ranging from 4 to 40 nM (Tsang et al., 2008), we might expect that the off-rate of Syk binding will vary in other cellular contexts. *In vitro* experiments have led to the description of Syk as a molecular switch that acts as an “OR” gate, whereby it is an equally active kinase when 1) engaged in ITAM binding in the open conformation *or* 2) when autophosphorylated (Tsang et al., 2008). However, our data here suggest that the two mechanisms are connected in a tightly controlled sequence. If Syk is bound sufficiently long to FcεRI to progress to a fully autophosphorylated state, it will remain active after dissociation into the cytosol. If the lifetime of ITAM binding is shorter, as with the Y130E mutant, both arms of the “OR” gate would be affected, leading to less efficient signaling. It follows that the lifetime of active, unbound Syk will also have a limited lifetime, because it will be reversed upon encounter with a phosphatase. Importantly, our results demonstrate that kinetic discrimination occurs at the level of Syk, where the extent of Syk phosphorylation at specific sites directs the cellular outcome.

We have demonstrated that Syk activation occurs through transient association with FcεRI. Live cell studies of Zap70 interacting with TCR have shown a transient interaction between kinase and receptor, with a range of dwell times from 0.2 s to tens of seconds (Bunnell et al., 2002; Klammt et al., 2015; O’Donoghue et al., 2013; Park et al., 2016). Although our measurements also found Syk binding to be transient, with lifetimes in the range of those reported for Zap70, it is interesting to note several significant differences between these two family members. Lillemeier and colleagues have used FRAP to examine the interaction time of Zap70 with TCR microclusters (Katz et al., 2017; Klammt et al., 2015). They found that I-B domain residues control the rates of transition between open and closed conformations of Zap70 and that Zap70 closed conformation mutants exhibited shorter TCR microcluster dwell times than wild type (Klammt et al., 2015). This shorter dwell time was associated with reduced phosphorylation of Zap70 by its Src family kinase, Lck (Klammt et al., 2015). For Syk, we found that shorter dwells times are not associated with impaired Lyn-dependent Syk phosphorylation; instead, Syk-Syk trans-phosphorylation is reduced. Recently, the same group examined mutations in the Zap70 I-A domain at Y126, the site analogous to Y130 in Syk (Katz et al., 2017). Zap70-Y126E dwell time was found to be similar to wild type, while Zap70-Y126F displayed longer dwell times. Cells expressing Zap70-Y126F showed impaired calcium signaling and reduced LAT and ERK phosphorylation, outcomes that are more similar to Syk-Y130E in mast cells. Therefore, while Syk and Zap70 share many structural and functional similarities, their distinctive characteristics underscore the need to avoid generalizations about their regulation. Another interesting distinction is the role of Syk in non-immune cells, where the Syk-Y130E substitution is described as a gain-of-function mutation since it promotes increased adhesion and microtubule dynamics (Yu et al., 2015). Clearly, the cellular context must be considered when dissecting the molecular mechanisms that govern protein function.

In summary, single molecule imaging has allowed us to quantify the transient interactions between FcεRI and Syk that drive mast cell signaling. We found that FcεRI-Syk binding dynamics are invariant with receptor aggregate size or receptor mobility. We also found that small changes in the dynamics of Syk binding, induced by the phosphomimetic mutation Y130E, are associated with significant alterations in the Syk phosphorylation profile and impaired mast cell responses. Taken together, these results support a model consistent with Syk-ITAM interactions that are independent of local receptor density, but highly sensitive to interaction time.

## Author Contributions

S.L.S and C.C. performed experiments. G.P. assisted with super-resolution experiments. P.R. and K.A.L. developed single particle tracking analysis. S.L.S, M.J.O. and K.A.L. developed off-rate analysis. C.C. generated CRISPR/Cas-9 Syk-KO cells. S.L.S carried out all single molecule data acquisition and analysis. D.S.L, B.S.W, W.S.H, K.A.L, C.C. and S.L.S designed and interpreted experiments. S.L.S, B.S.W and D.S.L wrote the manuscript with input from all the authors. D.S.L. directed the project.

## Acknowledgements

This work was supported by NIH R01GM100114 and the New Mexico Spatiotemporal Modeling Center (NIH) P50GM085273. We thank Ryan Suderman and Emanuel Salazar-Cavazos for useful discussions, Eunice Choi and Shayna Lucero for assistance with cell culture and degranulation assays, Dr. Jennifer Gillette for use of the Accuri C6 flow cytometer and Dr. Fu-Sen Liang for use of the mNG-PYL construct. We gratefully acknowledge use of the University of New Mexico (UNM) Cancer Center microscopy and flow cytometry facilities, as well as the NIH P30CA118100 support for these cores. The authors declare no competing financial interests.

## Experimental Methods

### Antibodies, Antigens, Reagents

All IgE used in this study was H1-DNP-ε-206 IgE, prepared as described previously (Liu et al., 1980). DNP-BSA containing ~15-25 DNP per BSA was purchased from Thermo Fisher Scientific (A23018) and DNP-lysine was purchased from Sigma (Nε-DNP-L-lysine hydrochloride, D0380). AF647-IgE was generated using NHS-chemistry as previously described (Schwartz et al., 2015), resulting in an average dye to protein ration of 3:1. Briefly, AF647-NHS ester dye (Thermo Fisher Scientific, A20006) was reacted at a 10:1 ratio with IgE, and excess dye was purified using a PD MidiTrap G-10 Sephadex desalting column (GE Healthcare, 28-9180-11). All references to pY Syk are named according to tyrosine positions in rodent Syk. Antibodies used are listed as follows: HRP conjugated anti-pY99 and anti-pY20 (Santa Cruz sc7020, sc508), anti-Mouse and anti-Rabbit secondary antibodies (Santa Cruz, SC-7020 SC-2004), Syk (Cell Signaling, D3Z1 XPR #13198), pY-Syk pY342 (Abcam, 195700), pY519/520 (Cell Signaling, C87C1, 2710), pY317 (Abcam, AB63515), pY346 (Cell Signaling, 2701), pY-Vav1 (Abcam, ab47282), Vav1 (Cell Signaling, 2502), LAT (LS- C46104), ERK (Cell Signaling, 137F5), pY-ERK (Cell Signaling, D13.14.4E). Dasatinib was purchased from Santa Cruz (SC-358114) and diluted in DMSO before further dilution to ensure solubility. Ionomycin and Phorbol ester 12-myristate 13-acetate (PMA) were from Sigma (I3909, 79326). Fura-2 AM and AF647-Phalloidin were from Thermo Fisher Scientific (A22287, F1201).

### Genome editing: Syk-KO Cell Line

Cas9-mediated DNA cleavage was used to knock-out (KO) the endogenous gene coding for the Syk protein in RBL-2H3 cells *via* the insertion of a premature stop codon in the first exon of the gene. A highly specific gRNA (5’-GGCCAGAGCCGCAATTACCT-3’) targeting the first exon of rat *Syk* was designed using the http://crispr.mit.edu/ portal and then sub-cloned into PX458 vector (Addgene plasmid #48138) for simultaneous expression of the gRNA, WT Cas9 and a GFP reporter. For the gRNA sub-cloning, two partially complementary oligonucleotides (Integrated DNA Technologies) were assembled by PCR. Gel purified PCR products were cloned into BbsI-digested PX458 using Gibson Assembly (NEB) following the manufacturer’s specifications. After cloning and sequencing, the final plasmid was used to transiently RBL-2H3 cells using the Amaxa system (Lonza) following the manufacturer’s recommendations. Positive, GFP-expressing, cells were selected by flow cytometry using an iCyt cell sorter and immediately platted at sub-optimal concentration in 96-well plates. Sub-clones were screened using western blotting to identify clones with no Syk expression. The absence of residual GFP expression in Syk KO clones was assessed using a Nikon TE2000 epi-fluorescence microscope.

### Constructs

Syk^mNG^-WT, Syk^mNG^-Y130E, Syk^mNG^-Y130F all refer to murine Syk DNA c-terminally fused to mNG via a short V5 linker (GGTAAGCCTATCCCTAACCCTCTCCTCGGTCTCGATTCTACG) tag. The V5 tag was added to mNG (Allele Biotechnology User License) via fusion PCR. Murine Syk cDNA (Thermo Scientific, MMM1013-202858457) fused to V5-mNG was generated by gene fusion PCR (Ho et al., 1989) using pfu ultra DNA polymerase (Stratagene). Y130E and Y130F mutations were introduced during gene fusion via primers with the corresponding mutation at codon 130. Total cDNA was amplified by PCR before subcloning into pcDNA3.1 directional topo vector (Life Technologies). All constructs were checked by sequencing.

### Cell Lines

RBL-2H3 cells (Metzger et al., 1986; Wilson et al., 2000) were cultured in MEM supplemented with 10% heat inactivated FBS, Puromycin and L-glutamine. To ensure authenticity, cells were checked for IgE binding and degranulation response after each thaw and used for up to 10 passages. Transfections were performed using the Amaxa system (Lonza) with Solution L and Program T-20. Stable cell lines were generated through G418 selection over a one week period followed by isolation of positive, mNG-expressing cells using an iCyt cell sorter with a 525/50 nm emission filter. Cell lines were checked for equal expression levels before experiments using an Accuri C6 flow cytometer (FL-1) and sorted again when needed. For all microscopy experiments, cells were plated into 8-well Lab Tek (Nunc) chambers at a density of 50,000 cell/well, primed overnight with unlabeled or AF647-IgE as indicated and then imaged within 24 hrs.

### Immunoblotting and Immunoprecipitation

3x10^6^ cells were plated on 100 mm tissue grade culture plates and primed overnight with IgE. Cells were stimulated with 0.1 μg/ml DNP-BSA or mock for indicated time at 37°C in HANKS buffered saline. Cell lysates from each plate were prepared in cold NP-40 lysis buffer. Protein concentration of cleared lysates was measured by bicinchoninic acid (BCA) assay (Pierce, Rockford, IL), and equal amounts of total protein were separated on a 4–15% polyacrylamide gel (Bio-Rad, Hercules, CA), transferred to nitrocellulose (iBlot transfer system; Life Technologies), probed with the indicated antibody according to manufacturer’s recommendation, and imaged using enhanced chemiluminescence on a Bio-Rad ChemiDoc. For detection of LAT phosphorylation, samples were immunoprecipitated using protein A-beads (Amersham GE Healthcare) loaded with anti-LAT primary antibody, then immune complexes were denatured and immunoblotted as described. Band intensities were quantified using the Biorad Image Lab Software (ImageLab 4.0.1) automatic band detection or via the volume tool with automatic background adjustment. Phosphorylated band intensity was normalized to total protein and all bands were adjusted for loading using beta-actin.

### Kinase Activity Assay

Syk-KO cells stably expressing Syk^mNG^-WT, Syk^mNG^-Y130E, Syk^mNG^-Y130F or mock were lysed and immune-precipitated using mNeonGreen nAb Agarose beads (Allele Biotechnology, ABP-NAB-MNGA050) according to manufacturer’s protocol. Kinase activity was assessed as described previously (Steinkamp et al., 2014). Syk protein concentration across precipitate samples was semi-quantified by SDS-PAGE followed by immunoblotting using an anti-Syk antibody. Blots were incubated, exposed and quantified as described in Western Blot methods section. Absorbance for Syk^mNG^-WT, Syk^mNG^-Y130E, Syk^mNG^-Y130F precipitates was normalized by relative protein levels with the absorbance for Syk-KO precipitates used as a baseline offset.

### Degranulation

Cells were grown in 24 well tissue culture plates for 24 hrs and primed with IgE. Cells were washed and stimulated in HANKS buffer with indicated concentration of DNP-BSA for 30 min at 37°C. Release of granular content was measured by β-hexosaminidase concentration as previously described (Schwartz et al., 2015). To determine calcium independent secretion potential, cells were treated with 1 µM Ionomycin (Sigma I3909-1mL) and 50 nM PMA (Phorbol 12-myristate 13-acetate, Sigma #P1585-1MG) for 30 min at 37°C.

### Calcium Imaging and Analysis

Measurements were carried out as previously described (Schwartz et al., 2015). The ratio of fluorescence intensity at 350 nm/380 nm excitation was calculated for each cell over time after background subtraction. Calcium ratio time courses were fit to a model:

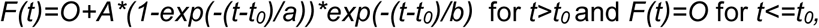

where *O* is the pre-stimulation signal, *t_0_* is the lag time between stimulation and response, *A* is the maximum rise height, and *a* and *b* describe the rise and decay kinetics respectively.

### Cytokine Release Measurements

Cells were primed with IgE and grown in 6 well tissue culture plates for 24 hr. Cells were washed and stimulated in HANKS buffer with 0.1 μg/mL DNP-BSA for 3 hrs. Supernatants were collected and mailed for shipment to RayBiotech, Inc according to company protocol. Cytokine quantification was performed in house by RayBiotech using their custom Rat Biomarker Quantibody^®^ platform. Samples were run in triplicate to assess technical variability. Results from at least three separate sample preparations for each cytokine were used to assess repeatability.

### Confocal Microscopy

Cells were primed with AF647-IgE and imaged using a Zeiss LSM800 laser scanning confocal microscope equipped with a 63× oil objective, and both 488 and 640 nm solid state diode laser excitation. Cells were imaged in HANKS buffer at 35°C. For photobleaching recovery studies, cells were stimulated for >3 min with 0.1 μg/ml DNP-BSA. Small regions of interest were identified and then a bleaching time series was acquired using the Zen2 software bleach ROI tool. For quantification, signal intensity recovery within the bleach regions was compared to signal in non-bleached regions to correct for photobleaching and background offset, then normalized to the pre-bleach intensity of the region. For actin labeling, cells were stimulated for 4 min with DNP-BSA before fixation with 4% PFA for 15 min followed by addition of approximately 6.6 μM AF647-phalloidin for 30 min.

### TIRF Microscopy Optical Setup

All TIRF imaging data was collected using an inverted microscope (IX71; Olympus) equipped with a 150× 1.45 NA oil-immersion, total internal reflection fluorescence objective (U-APO; Olympus). A 637 nm laser diode (HL63133DG, Thorlabs) was used for AF647 excitation, and a 488 nm laser (Cyan Scientific; Spectra-Physics) was used for fluorescence excitation. A quad-band dichroic and emission filter set (LF405/488/561/635-A; Semrock) was used for sample illumination and emission. Emission light was separated onto different quadrants of an EM-CCD camera (iXon 897; Andor Technologies), using a custom-built two-channel splitter with a 655 nm dichroic (Semrock) and 584/20 nm, 690/20 nm additional emission filters. Images had a pixel size of 0.106 µm and acquired at 10 frames per second (100 ms exposure time).

### Live Cell TIRF Microscopy

Samples were imaged in HANKS buffer at 35°C and temperature was maintained using a Bioptics Objective heater. Images were acquired before, during and for up to 5 min after addition of DNP-BSA at the indicated concentrations. To limit photobleaching of AF647-IgE, the 637 nm laser diode excitation was pulsed; every ten frames the laser was cycled on for two frames and then off for eight frames. A 488 nm laser power of 0.006 kW/cm^2^ was used for all imaging of single mNG-Syk molecules. If the mNG expression level in a cell of interest was too high, it was photobleached slightly out of TIRF with 0.38 kW/cm^2^ 488 nm laser power before image acquisition, meaning a population of dark/bleached Syk-^mNG^ also existed in all images. For Dasatinib treatment, cells were incubated with 1 μM Dasatinib for 30 min prior to imaging and also maintained during imaging and stimulation with DNP-BSA. For DNP-Lysine addition, cells were prepared and activated as described for 4-5 min allowing visible AF647-IgE aggregation, followed by addition of DNP-Lysine to a final concentration of 100 μM. Cells were imaged continually during this process.

### TIRF Microscopy Channel Registration

Quadrants of the EMCCD camera representing different fluorescence emission channels were aligned as follows. White light illumination through a channel alignment grid slide (Miraloma Tech) that contained a 20×20 array of 200 ± 50 nm holes at intra-hole distance (non-regular) of 3 ± 1 microns (total size ~60 x 60microns) was used to create an estimation of single point emitters appearing in both quadrants at regular samplings across the channel. The intensity passing through the holes was optimized to maximize number of photons without saturation. The localizations within each channel were then used to create a locally weighted transform matrix using the Matlab ‘fitgeotrans’ method with the ‘lwm’ option and the recommended twelve control points (Goshtasby, 1988).

### Image Masking

Intensity masking to quantify and compare the extent of IgE/Syk aggregation was carried out using image processing functions from Matlab (The MathWorks, Inc.) and the freely available Matlab package DipImage (Delft University of Technology). A whole cell mask was identified by Guassian filtering the image (kernel size = 2) and then thresholding using Matlab’s ‘multithresh’ function, followed by the DipImage ‘closing’ function. The overall image intensity was corrected for photobleaching by normalizing to the relative change in intensity within the whole cell mask over time. To identify aggregates within this cell mask, a two-step image segmentation process was used. (I) Images were smoothed and filtered using the DipImage ‘smooth’ and ‘dcc’ functions, then a threshold was identified using the Matlab ‘multithresh’ function and applied to the image, generating a mask representing regions of aggregation within the image. (II) To better isolate individual aggregates, a watershed transform was applied using the DipImage ‘watershed’ method with connectivity = 1, creating a second mask representing local areas of minimum intensities (watershed lines). The watershed lines from this step were then removed from the aggregate mask found in the first step to generate a final mask. The fluorescence intensity within each isolated mask was measured using the DipImage ‘label’ and ‘measure’ functions. The dynamics of aggregated AF647-IgE are relatively slow, so for analysis purposes, masks and corresponding mask intensities during ‘laser off’ frames between 637 nm laser excitation pulses (as described in the Live Cell TIRF methods section) were assumed to be static.

### Super-resolution imaging

Parental RBL cells labeled overnight with AF647-IgE (as described in Cell Lines method section) were washed three times with warm HANKS buffer to remove any unbound AF647-IgE, followed by stimulation at the indicated concentration of DNP-BSA in HANKS buffer for 5 min at 37°C. Cells were then quickly fixed with 4% PFA, 0.2% Gluteraldehyde for 2 hrs. Cells were extensively washed with PBS and once with 10 mM Tris-HCl (pH 7.2) for 10 min to quench reactive cross-linkers. Samples were imaged as previously described (Valley et al., 2015; van den Dries et al., 2013) using the same optical setup as described in the TIRF optical setup methods section and 637 nm laser power of ~1.7 kW/cm2. Images were acquired at 57 frames/s in TIRF. Between 10,000–20,000 frames were collected for each image reconstruction. The sample chamber was mounted in a three-dimensional piezostage (Nano-LPS; Mad City Labs, Madison, WI) with a resolution along the *xyz*-axes of 0.2 nm. Sample drift was corrected for throughout the imaging procedure using a custom-built stage stabilization routine.

### Super-resolution Image Reconstruction and Cluster Analysis

dSTORM images were analyzed and reconstructed with custom-built MATLAB functions as described previously (van den Dries et al., 2013; Yan et al., 2014). Images were reconstructed from between 5×10^5^ −10^6^ fit positions. To characterize the degree AF647-IgE-FcεRI aggregation, an implementation of Marlab’s Hierarchical clustering algorithm ‘linkage()’ using the ‘single’ method was used to group localizations into clusters. Code for this analysis, ‘Clustering Classes Version 2’, along with more information regarding the technique (Lin et al., 2016) is available through the STMC website [http://stmc.health.unm.edu/tools-and-data/].

### Diffusion Coefficient Estimation

Trajectories of single Syk^mNG^ (300 – 3000, see Table 1) were used for each indicated concentration of DNP-BSA to calculate a diffusion coefficient using a previously developed maximum likelihood estimation algorithm (Relich et al., 2016). 95% confidence intervals were calculated using a log likelihood ratio test (Pawitan et al., 2001).

### Code Availability

All computer code is available through request.

### Extracting Dissociation Rates from Single Particle Tracking of Low Signal to Noise Data

As has been nicely summarized previously (O’Donoghue et al., 2013; Presman et al., 2017; Woody et al., 2016), successful characterization of binding lifetimes from single molecule fluorescence data is highly dependent on the ability to (1) minimizing photobleaching while maintaining an adequate signal to noise ratio, (2) accurately localize single molecules and connect them across frames to build trajectories, and (3) accurately fit the distribution of trajectory lengths (representing the binding duration) to a binding model. Use of the mNeonGreen fluorophore (Shaner et al., 2013) allowed us to tune our laser power such that the rate of photobleaching was over an order of magnitude slower than our observed dissociation rates (Fig. S4A). The next sections detail our localization, tracking and off-rate parameter estimation.

### Localization Intensity-based Change Point Detection and Thresholding

We found that many Syk^mNG^ trajectories represented more than single molecule in a diffraction-limited area. We used an intensity change point algorithm (Ensign and Pande, 2009; O’Donoghue et al., 2013) to identify changes in intensity over the duration of the trajectory (Fig. S4B). This algorithm uses a Bayesian model selection technique to identify discrete changes of mean intensity for sequences of Poisson distributed data. The only parameter for the model is a Bayes factor used in a recursive decision procedure to divide each trajectory into segments of constant mean intensity. Similar change point profiles were found over a wide range of Bayes factor, but given the experimental variability we chose a conservative Bayes factor = 2.35 × 10^17^. We also characterized the expected fluorescence for single mNG molecules in TIRF using our experimental setup. Due to variability in fluorescence illumination and cell membrane morphology, we do not expect the fluorophore intensity to be exactly Poisson distributed. Therefore, we chose to experimentally characterize the fluorescent intensity distribution for single molecule mNG using an mNG fused to an unrelated, artificial PYL tag (mNG-PYL) (generously provided by Dr. Fu-Sen Liang, University of New Mexico) that exhibited significant non-specific binding at the membrane. Based on this empirical intensity distribution, we found <1% probability that a single mNG has intensity I_max_ ≥ 200 photons. I_max_ was therefore, used as a cutoff to exclude any localizations within our data that were obviously too bright to represent a single mNG. Any trajectories containing more than 2 localizations above I_max_ were not included in the analysis. We also found the mean fluorescence intensity for all mNG-PYL localizations I_mean_ = 49.1 photons. This value is consistent with the approximate size of the change point increments found using our change point analysis even though the analysis does not have any *a priori* knowledge of the intensity change magnitudes.

### Single Particle Tracking Method for Low Signal to Noise Data

#### (1) Segmentation of Particle Candidates

The segmentation and localization of particle candidates is handled through custom software. Segmentation is preformed by identifying potential emitters as the local maxima of a filtered image. We use a filter that convolves the image with the second derivative (Laplacian) of the Gaussian approximation to the microscope PSF. Candidate maxima are then thresholded based on their intensity in the filtered image.

Our localization procedure uses a new likelihood-based method. Like other established localization methods, we assume a 2D pixelated Gaussian PSF model under Poisson noise. Unlike the maximum likelihood estimation (MLE) approach employed in previous methods (Smith et al., 2010), we take a Bayesian approach based on finding the maximum a posteriori (MAP) estimate. The MAP parameter estimator uses a prior probability to regularize the likelihood function, making fast, but fallible local-optimization procedures such as the Newton-type methods more robust when used on data from high-speed low-intensity applications. The MAP estimator uses numerical optimization to find the parameters 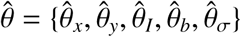, such that

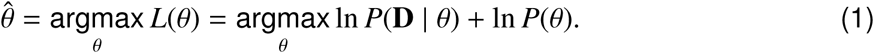

Here, *θ_x_* and *θ_y_* are the emitter’s position; *θ_I_* is the emitter intensity in photons; θ*_b_* is the mean background signal per pixel in photons; and *θ*_σ_ is the standard deviation of the apparent Gaussian PSF. The data is the image, **D**, a vector giving the photon counts at each pixel. The likelihood function, *P*(**D** |*θ*), under the Poisson noise assumption is described in (Smith et al., 2010). Finally, for the prior *P*(*θ*), we use a diagonal (separable) probability distribution, tuned for the emitter intensity profiles of our experiment:

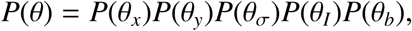

Where for an image of size (*s_x_, s_y_*),

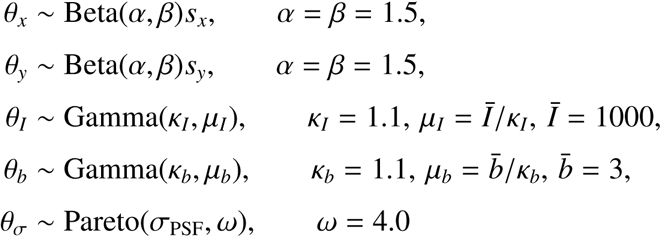

Our maximization procedure for the MAP estimation is performed with custom optimization software that uses a bound-constrained Newton’s method with reflective boundary conditions (Coleman et al., 1994), and deals with non-negative-definite Hessians using the modified Cholesky decomposition algorithm of Schnabel and Eskow (Schnabel and Eskow, 1999). For each iteration of the optimization procedure, we compute the objective function value *L*(θ) (Eq. 1), as well as the gradient ∇*L*(*θ*) and full Hessian matrix ∇^2^*L*(*θ*). Finally, we evaluate the quality of MAP estimates 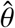, by assuring the Hessian 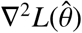 is negative-definite at the maxima and the gradient 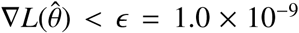. The observed Fisher information (Pawitan, 2001) at the maxima, 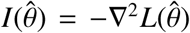, then can be used to establish the approximate shape of the posterior distribution *P*(*θ* | **D**) in the neighborhood of 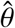, allowing accurate error bars on the estimated theta to be established.

Using the observed Fisher information, all emitter candidates that have estimates with a negative information value (which occurs on optimization failure) or those with very poor localization accuracy are discarded. The remaining candidates are then processed using the particle to trajectory assignment algorithm.

#### (2) Particle to Trajectory Assignments

Localized particles are processed into trajectories by creating connection hypotheses for particle assignments and finding the set of connection hypotheses that return the highest probability value. Inspired by previous approaches (Jaqaman et al., 2008), our method begins with a greedy frame-to-frame connection phase where particles are assigned to short trajectories by finding the most likely set of connections between particles localized in sequential frames. We take advantage of temporal symmetry by tracking particles both forwards and backwards in time, forming two sets of short trajectories. Particle connections that do not exist in both sets are omitted to reduce connection biases that ordinarily result from forward-time frame-to-frame tracking methods.

Next, the trajectory segments are connected over a sliding window in time, where only connections that occur between the starts or ends of trajectories found at the center of the sliding window are kept. This approach once again results in two lists of trajectory connections, depending on if the start or end of a short trajectory was found in the center of the sliding window. All connections that do not exist in both lists are once again removed to reduce the effect of connection bias. The processed trajectories are then scored based on the quality of their localizations. If a trajectory consists of mainly poor localizations, it will have a low score and thus be more likely to be removed. Any localizations that were not assigned to trajectories are discarded from the connection algorithm. Finally, the remaining trajectory segments are assigned to a global cost matrix (Jaqaman et al., 2008) in order to connect the gaps between short trajectories and more accurately reconstruct the trajectories of long lived binding events.

#### (3) Assignment Costs

From the list of trajectories, the time a trajectory starts (birth) and the time a trajectory ends (death) are collected. The probability of a trajectory beginning or ending on a particular frame is given by a user defined function of the minimum evidence value *e*_0_ = 0.05. The probability that two trajectories, *s_i_* and *s_j_*, should be connected is
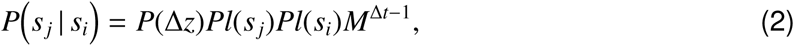

where Δ*_z_* represents the particle displacement that would be measured from connecting *s_j_* to *s_i_*. Here, *M* is a user defined probability of a missed localization in a trajectory and is only relevant when the difference in camera frame times between trajectories is greater than 1. The terms *Pl*(*s_i_*) and *Pl*(*s_j_*) represent the plausibility, defined as an upper bound of the probability, that *s_i_* and *s_j_* respectively represent a true particle. Particles are connected using a diffusion with drift model. The probability of two dimensional displacement, Δ*z* = [Δ*x*,Δ*y*], where the particle moves with diffusion constant *D* and drift rate **V** = [*V_x_*, *V_y_*] is
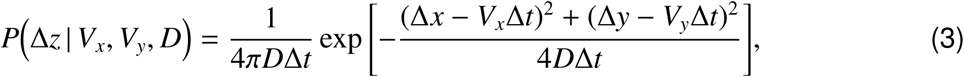

Here, Δ*t* is the number of camera frames elapsed during the measured displacement. The values for diffusion and drift are unknown and unique for every trajectory, so a normal distribution is used for the prior on drift constants and an inverse gamma distribution is used for the prior for diffusion constants:
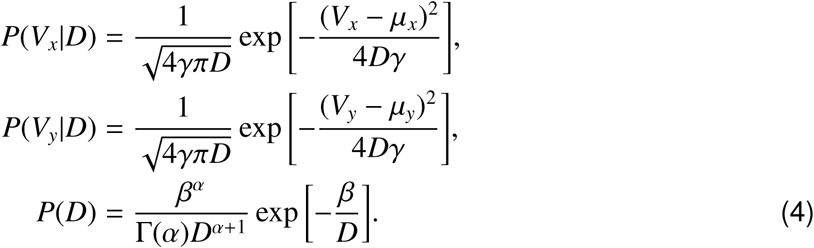

The hyperparameters *μ_x_*, *μ_y_*, and *γ* define the prior on drift velocities and the hyperparameters *α* and *β* define the prior on diffusion constants. The probability of a displacement is
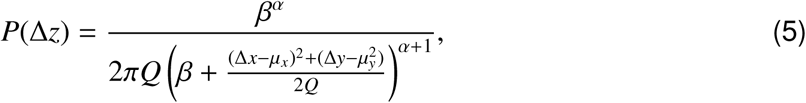

where *Q* = 2Δ*t*(1+*γ*Δ*t*).

The diffusion with drift hyperparameters are defined independently for each trajectory. Initially, a track uses default values, but as localizations are added, the values are updated utilizing the new information present in the additional displacement. If a trajectory’s motion changes over time, the hyperparameters are relaxed towards their default values to prevent over-fitting.

The plausibility terms are calculated as
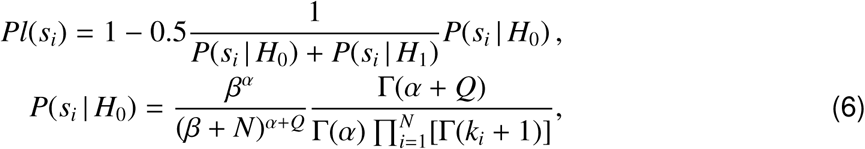

where *H*_0_ represents the null hypothesis that a localization was generated from a uniform background. The hypothesis *H*_1_ assumes the localization was generated by the usual Gaussian shaped point spread function. The hyperpriors on *H*_0_, *α* and *β*, are the same values used for estimating the background, *B*. For each subregion *H*_0_ is evaluated in, there are *N* pixels with *k* photon counts per pixel with a mean pixel count *Q*.

#### (4) Trajectory Filtering

Given a set of associated particles the probability that a trajectory is valid, *H*_1_, as opposed to invalid, *H*_0_, is 
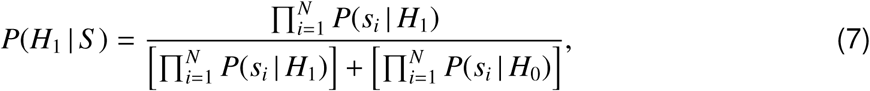

where *S* represents the vector of localizations associated to the trajectory under evaluation.

#### Off-rate Parameter Estimation

Observed trajectories of Syk^mNG^ at the plasma membrane represent individual molecules of Syk bound to FcεRI complexes. The simplest model that describes the unbinding of Syk from FcεRI:
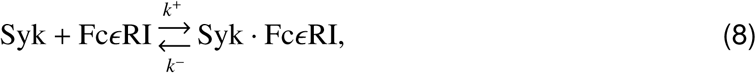

implies that for a Syk · FcεRI, the time until dissociation is described by random variable *τ* ~ Exp(*k*^−^). The exponential distribution has the memoryless property whereby if *τ* ~ Exp(*k*^−^), then *p*(*τ* = *t*) = *p*(*τ* = *t* + *t*_0_ | *τ* > *t*_0_), so that the distribution of *τ*, is unchanged even if the observation begins at arbitrary time *t*_0_ > 0 after the initial binding.

#### (I) The Geometric Distribution

The kinetics of Eq. 8 imply the continuous random variable *τ* is exponentially distributed; however, SPT trajectories are captured at a finite frame rate *t*_f_, so the durations *d* > 0 are measured in discrete units of frames. Hence, the distribution of trajectory durations in frames is described by a geometric distribution, *d* ~ Geo(*p*^−^). The geometric distribution is the discrete analog of the exponential, and the only discrete distribution with the memoryless property. The geometric distribution describes the number of trials until success for a repeated Bernoulli experiment with success probability 0 < *p* < 1 (e.g., the number of flips required to get heads using a *p*-weighted coin). In our context, a success represents unbinding which occurs with equal probability *p*− each frame. Thus, a trajectory is like a sequence of repeated coin flips, and the duration of a trajectory in frames represents the number of flips necessary to achieve success (unbinding). We can convert between *p*− and *k*− by taking advantage of the memoryless property of the exponential distribution. During any frame [*t*, *t* + *t*_f_), the probability of dissociation is independent of *t*:

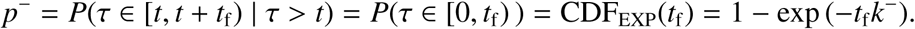

Hence the conversions are
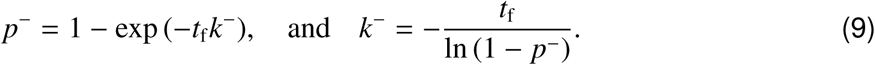

#### (II) Mixtures of Geometric Distributions

If the true kinetics of Syk unbinding is described by Eq. 8, then we should expect the durations 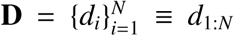 to follow a single geometric distribution
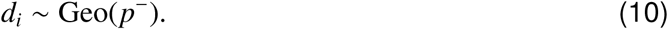

The single component geometric model is however a very simple model, and often more complicated models are necessary to adequately explain duration distributions. A chemically plausible extension assumes that there are different populations of bound Syk · FcεRI constructs that undergo dissociation with different rates
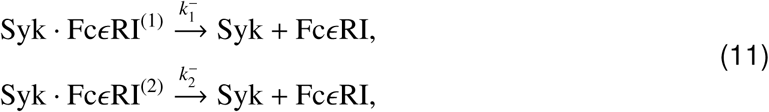

where 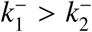. The proportion of Syk · FcεRI constructs in states 1 and 2 are represented by *α*_1_, *α*_2_ ∈ (0, 1) with *α*_1_ + *α*_2_ = 1. For the kinetic system of Eq. 11 the duration of a trajectory in frames is given by a 2-component mixture of geometric distributions
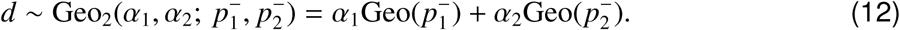

Extensions to *M* = 3 or more component geometric distributions are straight forward, where
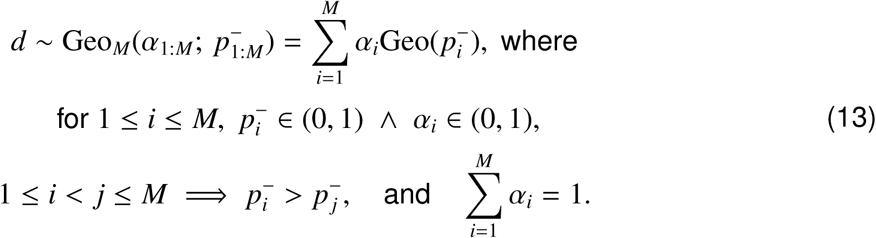

#### (III) Bayesian inference for geometric mixture models

Using the geometric mixture models (Eq. 13), there are two general classes of procedures we can perform: parameter estimation, and model selection. In parameter estimation we assume a model for the data *d_i_* ~ Geo*_M_*(*θ*) and estimate the most likely parameters *θ* for that model given the observed data **D**. In model selection we compare two or more models to determine which model best describes the observed data, after which, parameter estimation can be preformed on the most likely model. For both types of procedures we utilize the Bayesian approach (Murphy, 2012).

The Bayesian approach to parameter estimation begins with an application of Bayes’ rule to the likelihood for model 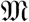,
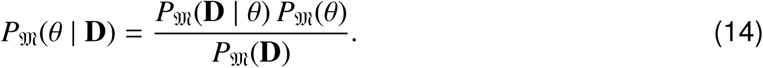

The posterior probability distribution 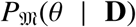, is the ultimate mathematical objective of Bayesian parameter estimation. It represents the distribution of the true parameters *θ* given the data **D**. We can compute the posterior using Eq. 14 to combine the *likelihood* 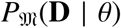 and the *prior* 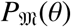. The marginal *likelihood* 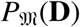, is simply a normalization constant that we will avoid computing in our MCMC approach.

(III.A) *The likelihood function*: For the *M*-component geometric mixture distribution (Eq. 13), the free parameters^1^ are 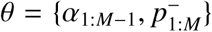, and the likelihood of an individual trajectory duration is 
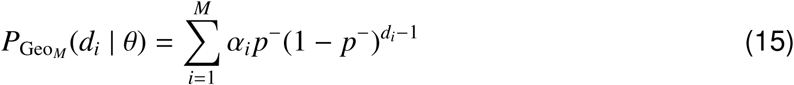

using Eq. 15 we define the log-likelihood function, parametrized on *θ*, assuming a fixed dataset

**D** = *d*_1:*N*_ of *N* independent trajectories,
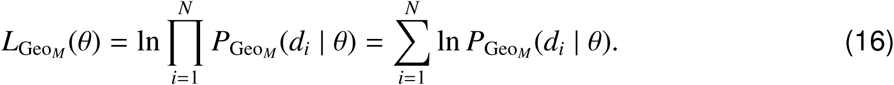

(III.B) *Priors*: To ensure our inferences are solely based on the data at hand, we assume no prior information on our parameters and use a uniform prior over (0, 1) for each *α_i_* and the Jeffrey’s prior (Murphy, 2012) for dissociation probabilities:
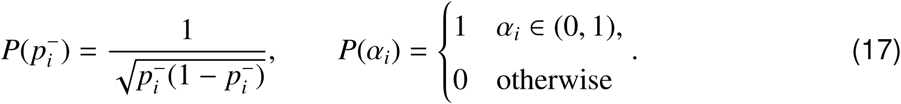

(III.C) *Extending models to multiple conditions*: In order to compare kinetic parameters across different experimental conditions, we can build a joint model over *K* > 1 conditions, where the data 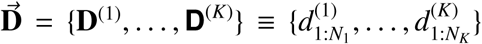, and each 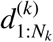 is a collection of the *N_k_* > 0 observed trajectories from experimental condition *k*.

A joint *K*-condition *M*-component geometric mixture model, 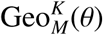, has *K*(2*M* – 1) free parameters after accounting for the constraints on *α* fractions, 
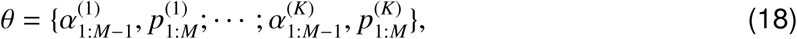

and the joint likelihood function is
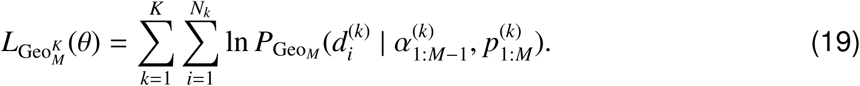

Furthermore, we can investigate restrictions of the *K*-condition *M*-component model by assuming all of the *K* conditions share one or more of the underlying rate or fraction parameters. For example, if there is reason to believe all conditions share the same fast rate, but may have different values for the slower rate, we can create a model where 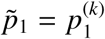 for each *k*, yet each condition retains it’s own 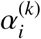 and 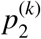 rates, so that we are left with a *K*(2*M* – 2) + 1 dimension model with parameters
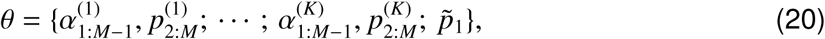

Similar restricted *K*-component models can be proposed by fixing any of the *α* or *p* parameters across the conditions.

#### (IV) Bayesian parameter estimation using Markov chain Monte Carlo

Given a model 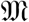 and data **D**, Eq. 14 shows the posterior probability of the parameters θ given the data is proportional to the likelihood times the prior,
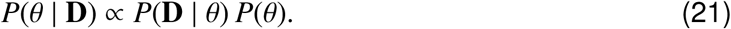

The Markov chain Monte Carlo method (MCMC) (Murphy, 2012) can be used to sample from the posterior distribution using the unnormalized formula of Eq. 21. This requires only computing the likelihood (Eq. 19), and the prior (Eq. 17), and does not require the marginal likelihood *P*(D).

To perform parameter inference on the models, we implemented a MCMC sampler for the parameter space of the general *K*-condition *M*-component geometric model (Eq. 18) and the related restricted models (e.g., Eq. 20). We use a 1D Gaussian random-walk proposal algorithm that changes a single dimension at each MC iteration, cycling through all parameters in sequence. Since all variables are constrained to the interval (0, 1), we used a 1D proposal Gaussian with *σ* = 0.05 to achieve adequate acceptance rates and mixing speed. Any proposal that was non-feasible because of violation of constraints (Eq. 13) is treated as a normal rejection step. This is equivalent to assigning zero likelihood to infeasible points. In general for MCMC runs we collect *S* = 10^6^ samples after rejecting the first *S*/4 samples as a burn-in.

If the true parameters are *θ*\*, the estimated theta 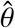 that minimizes the euclidean distance 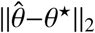 is the posterior mean, otherwise known as the minimum mean squared error estimator. The MCMC samples allow us to easily estimate the posterior mean and covariance with the sample mean and covariance. Indeed, nearly any probabilistic query or hypothesis that can be formulated about the posterior distribution can be estimated directly using the MCMC samples.

#### (V) Bayesian model selection using Markov chain Monte Carlo

Given dataset **D**, and a set of *J* > 1 models 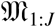, we can use Bayesian model selection to estimate the posterior likelihood of each model given the data 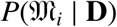. Bayes’ rule shows that,
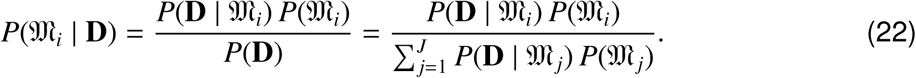

We assume a uniform prior over models 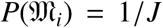, so in order to estimate the posterior distribution for the models with Eq. 22, we only need to estimate the model evidence,
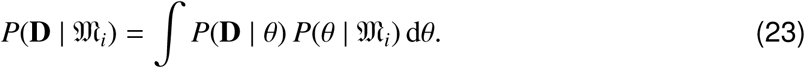

We use a Laplacian approximation (Tierney and Kadane, 1986) to estimate the value of the integral in Eq. 23. This method assumes the parameter posterior distribution 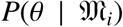 is approximately normal. Then the sample mean and covariance from an MCMC run are used to form a Gaussian approximation to the posterior. With 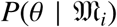 Gaussian, the integral of Eq. 23 is then in a form to which the Laplace method is applicable.

In practice, running an MCMC estimate of 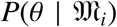 for each model 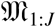, allows us to compute a posterior probability for the models 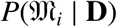, and the model with highest posterior probability is the maximum a posteriori (MAP) model estimate,
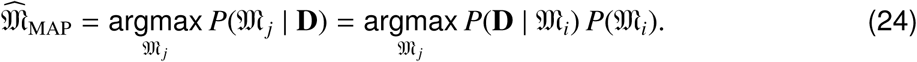

As a consequence of the MCMC sampling to estimate Eq. 22, after finding 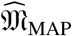 we already have the MCMC sample of the parameter posterior 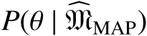.

#### (VI) Conditional likelihood for data with a minimum track length cutoff

Very short trajectories are sometimes false positives that result from spurious localizations or tracking artifacts. Longer trajectories tend to be less susceptible to such noise, hence we imposed a lower bound *m* ≥ 1 on track lengths that are used in rate model inferences. Therefore, given dataset **D** = *d*_1:*N*_, we know a priori that *d_i_* ≥ *m*, so we replace 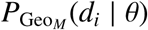 in the likelihood function of Eq. 16 with the conditional probability,
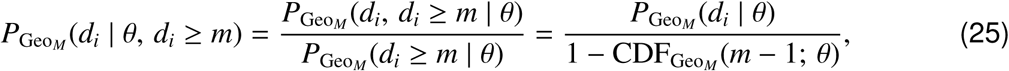

where for 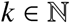,

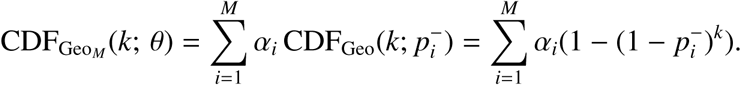

In the single component case (*M* = 1), the geometric model has the unique memoryless property which allows the simplification 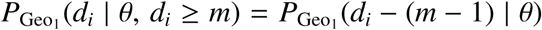, but for *M* = 2 or more component mixture models, the memoryless property does not hold, and Eq. 25 must be used for minimum track length thresholded datasets.

#### Model selection results

We used Bayesian inference methods to answer several questions about the kinetic behavior of Syk*^mNG^* across different DNP-BSA doses and between Syk*^mNG^* isoforms (Syk*^mNG^*-WT, Syk*^mNG^*-Y130E and Syk*^mNG^*-Y130F). For all conditions we used a minimum track length cutoff *m* = 6.

First, we compared *M* = 1, *M* = 2, and *M* = 3 component models for each condition independently. For all conditions (except with Dasatanib treatment) we found that the *M* = 2 model had a higher posterior probability compared to the *M* = 1 model, indicating that our data is clearly not well described by a single geometric distribution (Fig. S4C, left). While we did find that for some conditions (high doses for Syk*^mNG^*-WT and Syk*^mNG^*-Y1 30F), an *M* = 3 component model had a higher posterior probability, the relative improvement in the fit (Fig. S4C, right vs center) could not justify the additional complexity it added when considering cross-condition model comparisons, given the available number of trajectories. For the *M* = 2 component model, 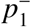 corresponds to the faster dissociation rate *k_f_* and 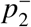 the slower rate *k_s_*. We found that parameter estimates for *k_f_* were similar across conditions when fit independently, while *α_s_* and *k_s_* varied over a wide range. We assume *k_f_* represents non-specific binding events that we don’t expect to vary across conditions.

Assuming the 2-component model with a *k_f_* shared across conditions, we next asked if there was evidence to support a model in which the slow rate *k_s_* is also shared. We compared a model in which a single *k_s_* was constrained to be shared across conditions, versus a model in which each condition had an independent *k_s_* rate. Across antigen dosage conditions we found a significantly higher posterior probability for a model in which both *k_f_* and *k_s_* are shared and only *α_s_* changes per condition. In contrast, across cell-line conditions, comparing Syk*^mNG^*-Y130E to either Syk*^mNG^*-WT or Syk*^mNG^*-Y130F, we found a significantly higher posterior probability for a model where the slow rate *k_s_* is variable.

In addition, we use the MCMC samples from the posterior distribution (Fig. S4D) to investigate the difference between the slow kinetic rates *k_s_* across different conditions. In Fig. S4E we plot the cumulative probability distribution for differences 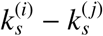 based on MCMC sample of the posterior of the geometric parameters 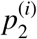 and 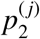 after conversion to kinetic rates (Eq. 9). These results show, the difference between *k_s_* values for WT and YF are almost certainly less than 0.1 s^−1^ and for WT versus YE almost certainly greater than 0.1 s^−1^ (Fig. S4E).

#### Single Molecule Syk^mNG^ Binding Simulations

To verify our analysis, Syk^mNG^ data was simulated using three populations of simulated emitters. The first population of emitters represented the background fluorescence described by a wide PSF sigma of 4 pixels and fast diffusion constant of 5 pixels^2^/frame. The other two populations represented Syk^mNG^ molecules with a PSF sigma of 1 pixel and a diffusion constant of 0 pixels^2^/frame or 0.5 pixels^2^/frame, matching out experimental observation. Simulation off rates for each population corresponded to the fit kf or ks values from our results (Table 1). An exponential rate was also used to draw the starting time for each emitter, reflective of the α_f_ and α_s_ parameter. A set number of emitters were drawn for each population at the beginning of the simulation. All emitters were simulated with a mean photon rate per frame matching our experimental observations, from which, emission times for each photon were drawn. An emitter was allowed to diffuse freely up until the time of photon emission, at which point the location of the captured photon was recorded as a random displacement from the actual emitter location drawn from a normal distribution scaled by the corresponding PSF sigma of the emitter. All photon locations were summed to create a Poisson corrupted image series of emitter dynamics that was evaluated using our tracking and track lifetime analysis. Binding parameters (k_f_,k_s_,(α_s_) found from analyzing these simulations were all within one standard error of the input parameters (Fig. S4F).

---

**Figure S1.**
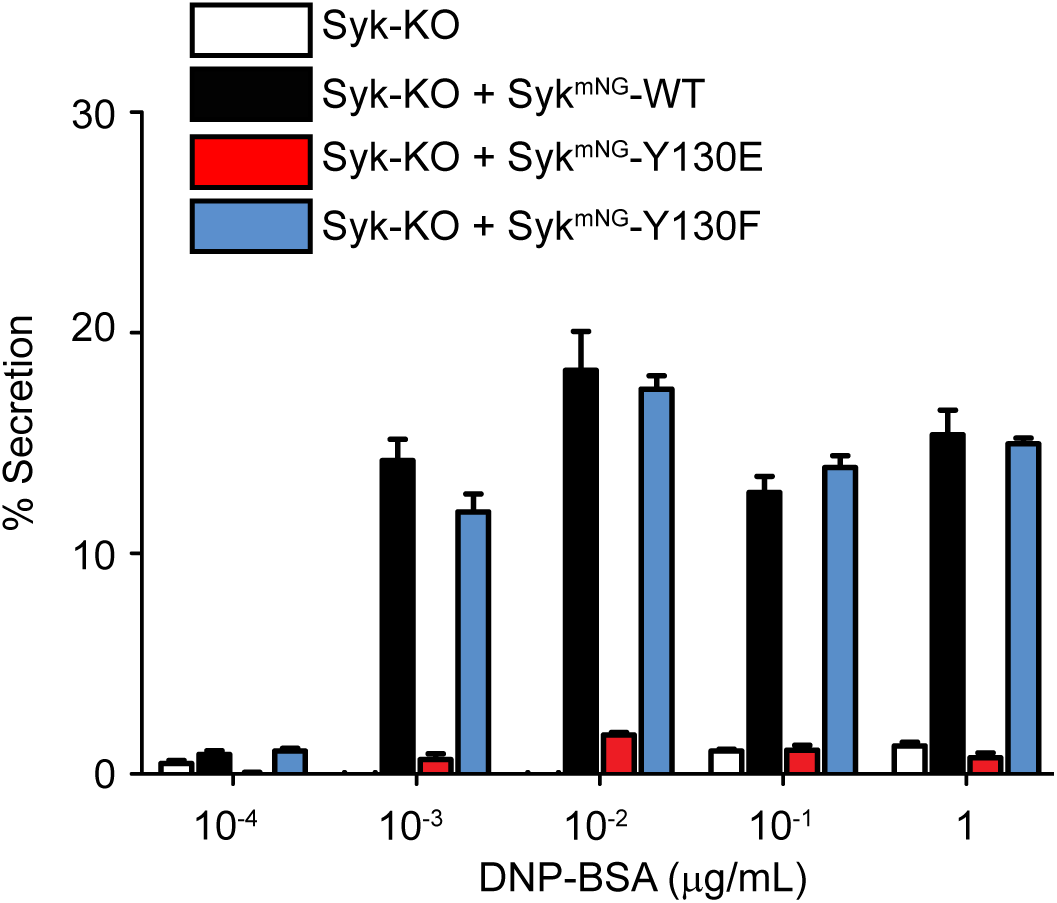
Mast cell degranulation response for Syk^mNG^-WT and mutants. Degranulation assay showing fraction of β-hexosaminidase released after 30 min of incubation over a range of DNP-BSA doses in Syk-KO cells (white bars) or in Syk-KO cells stably expressing Syk^mNG^-WT (black bars), Syk^mNG^-Y130E (red bars), Syk^mNG^-Y130F (blue bars).

**Figure S2.**
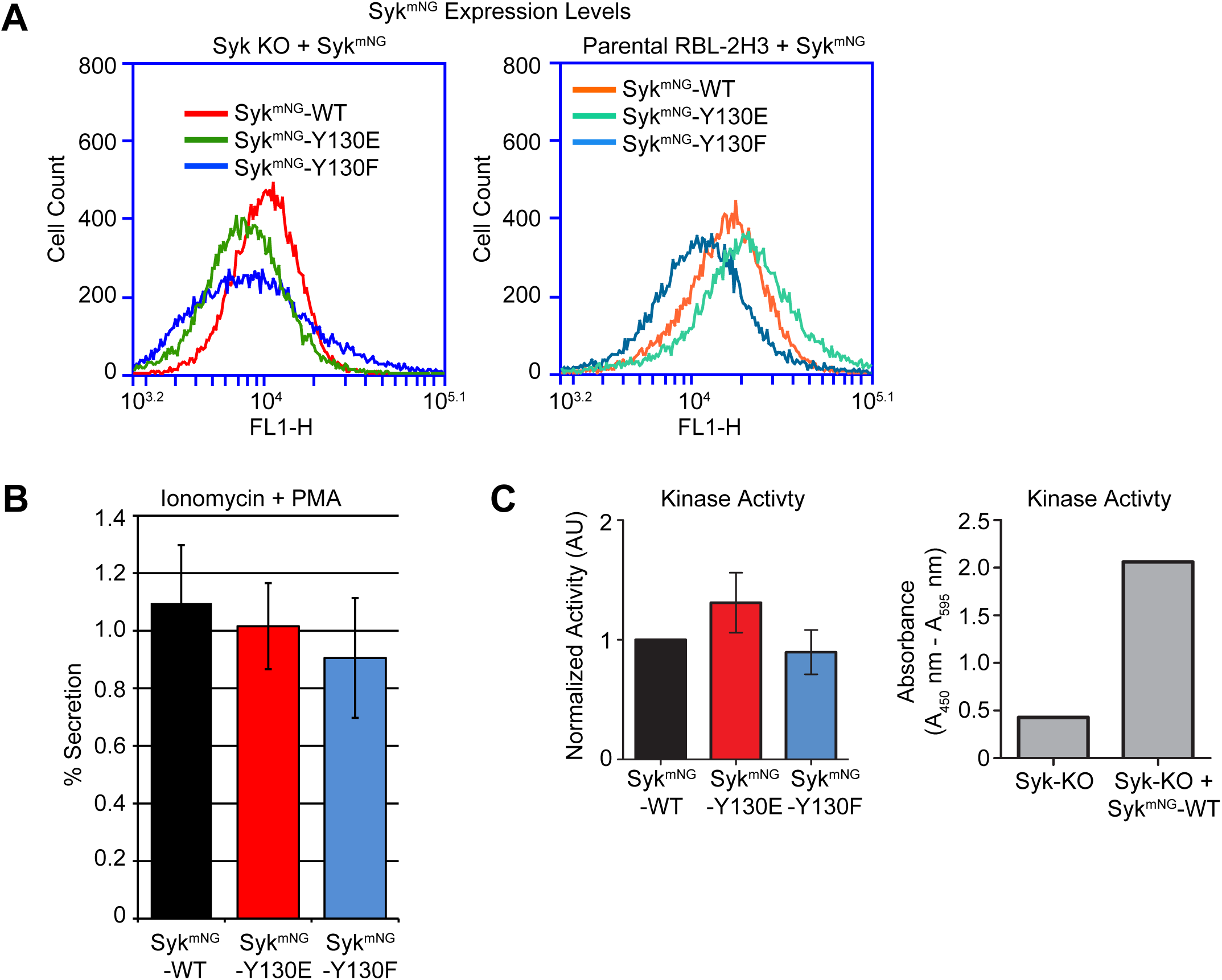
Characterization of stable Syk^mNG^ cell lines. (A) Syk^mNG^-WT, Syk^mNG^-Y130E, and Syk^mNG^-Y130F expression quantified using flow cytometry in Syk-KO or parental RBL-2H3 cell lines. (B) β-hexosaminidase release after treatment with Ionomycin and PMA in Syk-KO cell lines expressing Syk^mNG^-WT, Syk^mNG^-Y130E, or Syk^mNG^-Y130F. (C) Left: Relative kinase activity for Syk-KO cell lines expressing Syk^mNG^-WT, Syk^mNG^-Y130E, or Syk^mNG^-Y130F. Values normalized to Syk^mNG^-WT for comparison. Right: Raw absorbance measurements in Syk-KO cells with (positive control) or without (negative control) expression of Syk^mNG^-WT.

**Figure S3.**
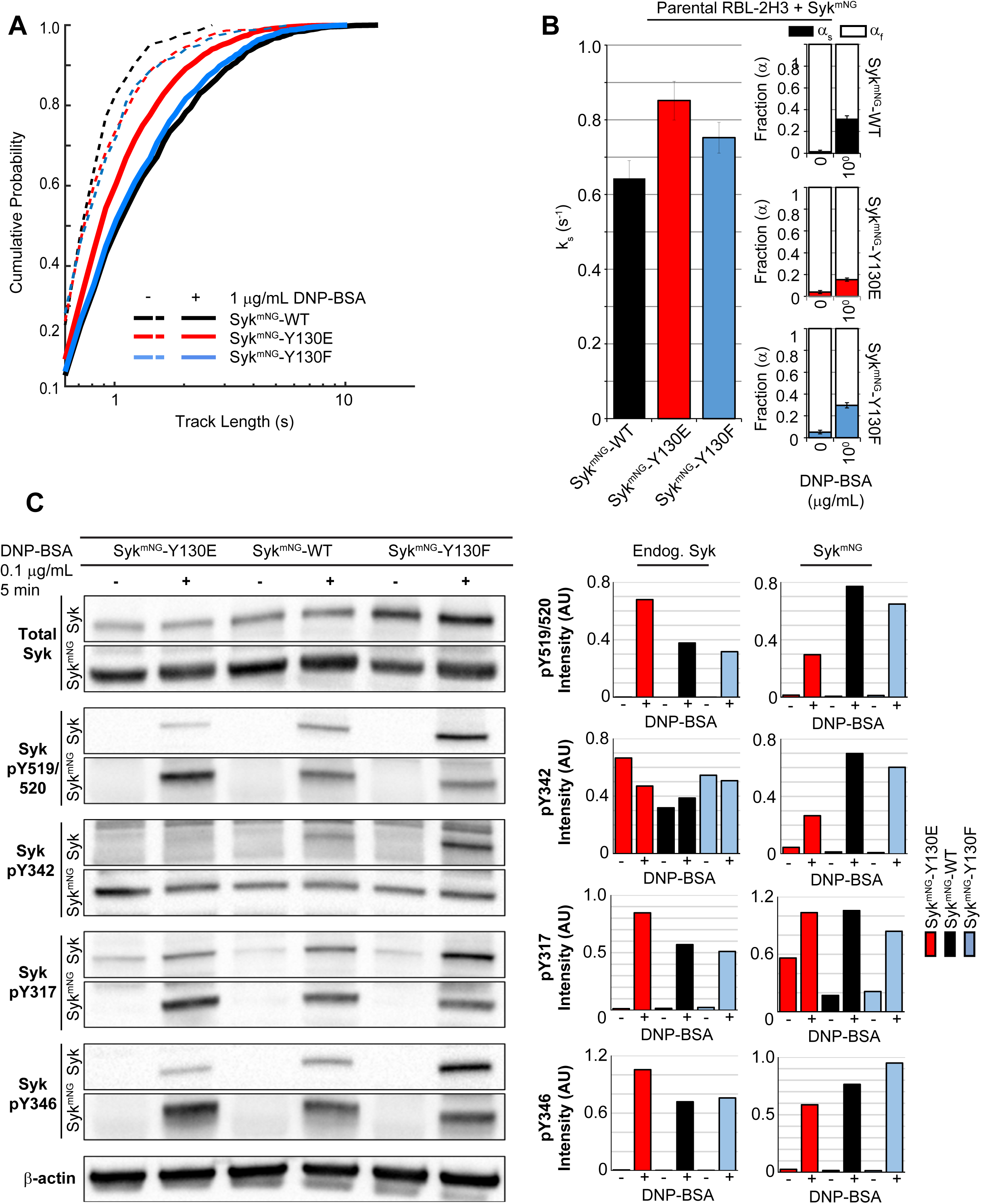
Syk^mNG^ -Y130E exhibits a faster off-rate and altered phosphorylation in the presence of endogenous Syk within parental RBL-2H3 cells. (A) Cumulative probability distributions for Syk^mNG^ trajectory lengths in parental RBL-2H3 cells expressing Syk^mNG^-WT, Syk^mNG^-Y130E, or Syk^mNG^-Y130F both before (dashed lines) and after (solid lines) addition of 1 μg/mL DNP-BSA. (B) Slow off-rate value (k_s_) found when fitting distriubtions in (A). Fraction of slow off-rate component (α_s_) increases with DNP-BSA dose for Syk^mNG^-WT and each mutant (right). Error bars are a 68% credible interval as described in Methods. (C) Western blot detection of Syk phosphorylation profile in parental RBL-2H3 cells expressing Syk^mNG^-WT, Syk^mNG^-Y130E, or Syk^mNG^-Y130F in response to stimulation with 0.1 μg/mL DNP-BSA for 5 min. Blots show both endogenous Syk phosphorylation (top) and Syk^mNG^ phosphorylation (bottom) at each site for all three cell types. Quantification of Syk phosphorylation from one representative experiment; endogenous Syk phosphorylation (left) and Syk^mNG^ phosphorylation (right).

**Figure S4.**
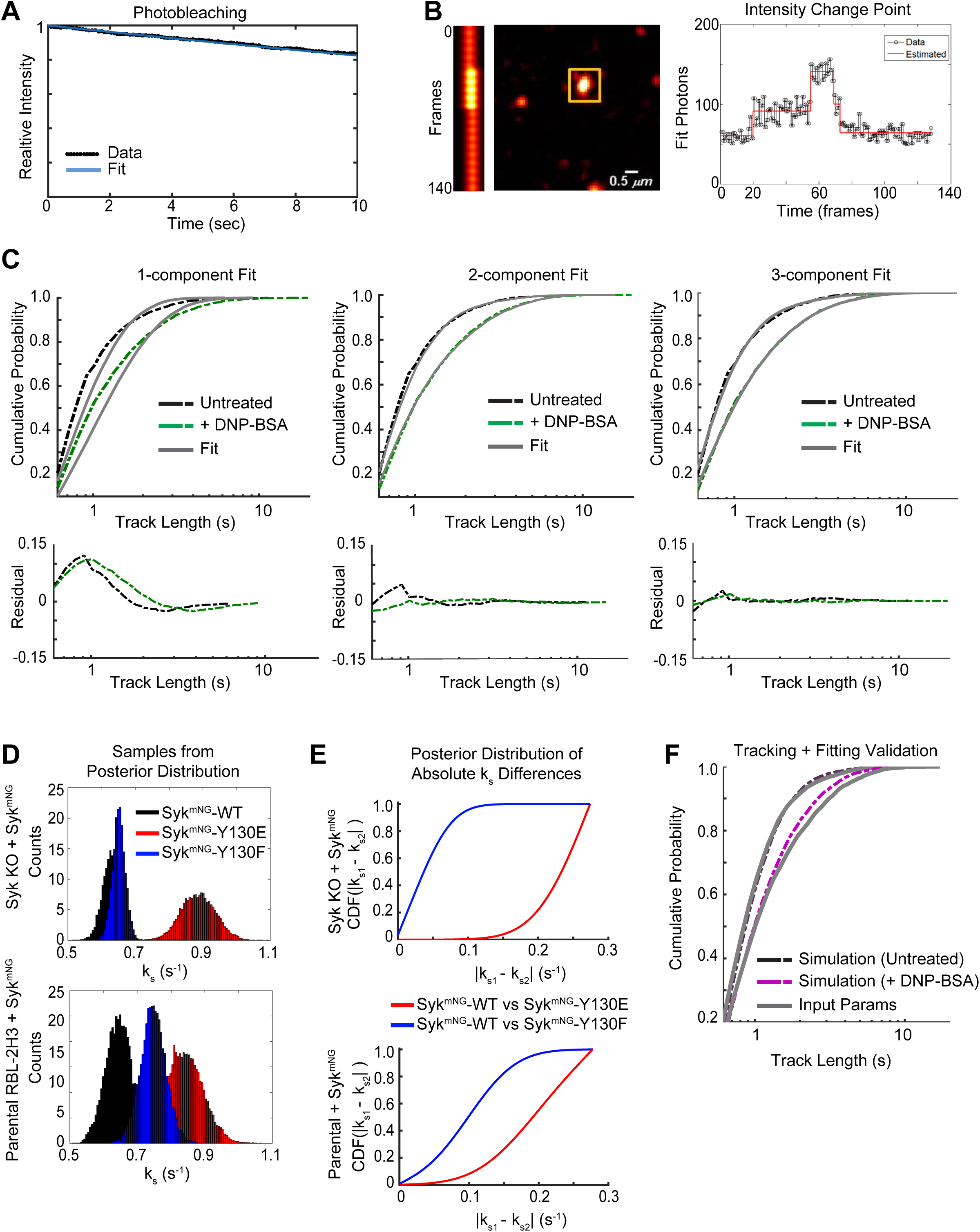
Overview of Syk trajectory off-rate fitting approach. (A) The photobleaching rate for mNG under the experimental imaging setup used for all single molecule data collection. The intensity of the whole cell was monitored under continuous illumination over time. A rate of 0.0193 s^−1^ with 95% confidence interval (0.01928, 0.01933) and a goodness of fit R-square value = 0.9994 was found using the Matlab Curve Fitting Tool to fit the relative intensity to a custom equation f(x) = exp(-a*x). (B) Example intensity profile and change point analysis for a single Syk^mNG^ trajectory. (Left) Kymograph of particle intensities over time. Scale bar, 0.5 µm. (Right) Intensity profile and resulting identified change points using change point analysis. (C) Comparison of fitting models for Syk^mNG^ trajectory off-rates. Cumulative probability distribution of Syk^mNG^ trajectory lengths before (black) and after (green) addition of 1 μg/mL DNP-BSA compared with fitting (grey) using a 1, 2 and 3 component geometric mixture model. Residuals of data and fit are shown below for each model. (D) The sampled posterior distribution of the k_s_ parameter value using Markov-Chain Monte Carlo for data from Syk^mNG^-WT, Syk^mNG^-Y130E, or Syk^mNG^-Y130F trajectories in Syk-KO cells (top) or parental RBL cells (bottom). (E) Cumulative probability distributions for expected differences in k_s_ parameter value between Syk^mNG^-WT and Syk^mNG^-Y130E (red) or Syk^mNG^-WT and Syk^mNG^-Y130F (blue) in Syk-KO cells (top) or parental RBL cells (bottom). See Methods for more details. (F) Validation of tracking and fitting approach. Parameters similar to those found for Syk^mNG^-WT in untreated and DNP-BSA stimulated cells were used to generate simulated image series of single molecule binding events that were then analyzed. Cumulative probability distribution of resulting trajectory lengths (untreated-black, DNP-BSA stimulated-magenta) are compared with the expected distribution given the input parameters (grey). Fit parameters found from the simulation are within one standard error of the input parameters. See Methods for more details.

## Footnotes

1 The constraints on *α* imply that *α_M_* is not free. Given *α*_1:*M*−1_ it has the fixed value 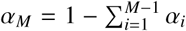, and so we treat models as parameterized on *α*_1:*M*−1_.

## References

Andrews, N.L., J.R. Pfeiffer, A.M. Martinez, D.M. Haaland, R.W. Davis, T. Kawakami, J.M. Oliver, B.S. Wilson, and D.S. Lidke. 2009. Small, Mobile FcεRI Receptor Aggregates Are Signaling Competent. Immunity. 31:469–479.

Arias-Palomo, E., M.A. Recuero-Checa, X.R. Bustelo, and LlorcaO. 2009. Conformational rearrangements upon Syk auto-phosphorylation. Biochimica et biophysica acta. 1794:1211–1217.

Au-Yeung, B.B., S. Deindl, L.Y. Hsu, E.H. Palacios, S.E. Levin, J. Kuriyan, and A. Weiss. 2009. The structure, regulation, and function of ZAP-70. Immunol Rev. 228:41–57.

Barker, S.A., K.K. Caldwell, A. Hall, A.M. Martinez, J.R. Pfeiffer, J.M. Oliver, and B.S. Wilson. 1995. Wortmannin blocks lipid and protein kinase activities associated with PI 3-kinase and inhibits a subset of responses induced by Fc epsilon R1 cross-linking. Mol Biol Cell. 6:1145–1158.

Barker, S.A., D. Lujan, and B.S. Wilson. 1999. Multiple roles for PI 3-kinase in the regulation of PLCgamma activity and Ca2+ mobilization in antigen-stimulated mast cells. J Leukoc Biol. 65:321–329.

Brdicka, T., T.A. Kadlecek, J.P. Roose, A.W. Pastuszak, and A. Weiss. 2005. Intramolecular regulatory switch in ZAP-70: analogy with receptor tyrosine kinases. Mol Cell Biol. 25:4924–4933.

Bunnell, S.C., D.I. Hong, J.R. Kardon, T. Yamazaki, C.J. McGlade, V.A. Barr, and L.E. Samelson. 2002. T cell receptor ligation induces the formation of dynamically regulated signaling assemblies. J Cell Biol. 158:1263–1275.

Carsetti, L., L. Laurenti, S. Gobessi, P.G. Longo, G. Leone, and D.G. Efremov. 2009. Phosphorylation of the activation loop tyrosines is required for sustained Syk signaling and growth factor-independent B-cell proliferation. Cell Signal. 21:1187–1194.

Chen, C.-H., V.A. Martin, N.M. Gorenstein, R.L. Geahlen, and C.B. Post. 2011. Two closely spaced tyrosines regulate NFAT signaling in B cells via Syk association with Vav. Molecular and cellular biology. 31:2984–2996.

Chu, D.H., H. Spits, J.F. Peyron, R.B. Rowley, J.B. Bolen, and A. Weiss. 1996. The Syk protein tyrosine kinase can function independently of CD45 or Lck in T cell antigen receptor signaling. EMBO J. 15:6251–6261.

Cocucci, E., F. Aguet, S. Boulant, and T. Kirchhausen. 2012. The first five seconds in the life of a clathrin-coated pit. Cell. 150:495–507.

Coleman, T.F., and Y. Li. 1994. On the convergence of interior-reflective Newton methods for nonlinear minimization subject to bounds. Mathematical Programming. 67:189–224.

Costello, P.S., M. Turner, A.E. Walters, C.N. Cunningham, P.H. Bauer, J. Downward, and V.L. Tybulewicz. 1996. Critical role for the tyrosine kinase Syk in signalling through the high affinity IgE receptor of mast cells. Oncogene. 13:2595–2605.

Das, J., M. Kardar, and A.K. Chakraborty. 2009. Positive feedback regulation results in spatial clustering and fast spreading of active signaling molecules on a cell membrane. J Chem Phys. 130:245102.

Deckert, M., S. Tartare-Deckert, C. Couture, T. Mustelin, and A. Altman. 1996. Functional and physical interactions of Syk family kinases with the Vav proto-oncogene product. Immunity. 5:591–604.

Deindl, S., T.A. Kadlecek, T. Brdicka, X. Cao, A. Weiss, and J. Kuriyan. 2007. Structural basis for the inhibition of tyrosine kinase activity of ZAP-70. Cell. 129:735–746.

Eiseman, E., and J.B. Bolen. 1992. Engagement of the high-affinity IgE receptor activates src protein-related tyrosine kinases. Nature. 355:78–80.

Ensign, D.L., and V.S. Pande. 2009. Bayesian single-exponential kinetics in single-molecule experiments and simulations. The journal of physical chemistry. B. 113:12410–12423.

Feng, C., and C.B. Post. 2015. Insights into the allosteric regulation of Syk association with receptor ITAM, a multi-state equilibrium. Physical chemistry chemical physics: PCCP.

Geahlen, R.L. 2009. Syk and pTyr’d: Signaling through the B cell antigen receptor. Biochimica et Biophysica Acta - Molecular Cell Research. 1793:1115–1127.

Goshtasby, A. 1988. Image registration by local approximation methods. Image Vision Comput. 6:255–261.

Graham, T.E., J.R. Pfeiffer, R.J. Lee, D.F. Kusewitt, A.M. Martinez, T. Foutz, B.S. Wilson, and J.M. Oliver. 1998. MEK and ERK activation in ras-disabled RBL-2H3 mast cells and novel roles for geranylgeranylated and farnesylated proteins in Fc epsilonRI-mediated signaling. J Immunol. 161:6733–6744.

Hlavacek, W.S., A. Redondo, H. Metzger, C. Wofsy, and B. Goldstein. 2001. Kinetic proofreading models for cell signaling predict ways to escape kinetic proofreading. Proc Natl Acad Sci U S A. 98:7295–7300.

Ho, S.N., H.D. Hunt, R.M. Horton, J.K. Pullen, and L.R. Pease. 1989. Site-directed mutagenesis by overlap extension using the polymerase chain reaction. Gene. 77:51–59.

Hopfield, J.J. 1974. Kinetic proofreading: a new mechanism for reducing errors in biosynthetic processes requiring high specificity. Proceedings of the National Academy of Sciences of the United States of America. 71:4135–4139.

Hutchcroft, J.E., R.L. Geahlen, G.G. Deanin, and J.M. Oliver. 1992. Fc epsilon RI-mediated tyrosine phosphorylation and activation of the 72-kDa protein-tyrosine kinase, PTK72, in RBL-2H3 rat tumor mast cells. Proc Natl Acad Sci U S A. 89:9107–9111.

Ichinose, J., M. Morimatsu, T. Yanagida, and Y. Sako. 2006. Covalent immobilization of epidermal growth factor molecules for single-molecule imaging analysis of intracellular signaling. Biomaterials. 27:3343–3350.

Isaacson, C. 1997. Syk Activation and Dissociation from the B-cell Antigen Receptor Is Mediated by Phosphorylation of Tyrosine 130. Journal of Biological Chemistry. 272:10377–10381.

Jaqaman, K., D. Loerke, M. Mettlen, H. Kuwata, S. Grinstein, S.L. Schmid, and G. Danuser. 2008. Robust single-particle tracking in live-cell time-lapse sequences. Nature methods. 5:695–702.

Johnson, S.a., C.M. Pleiman, L. Pao, J. Schneringer, K. Hippen, and J.C. Cambier. 1995. Phosphorylated immunoreceptor signaling motifs (ITAMs) exhibit unique abilities to bind and activate Lyn and Syk tyrosine kinases. Journal of immunology (Baltimore, Md.: 1950). 155:4596–4603.

Katz, Z.B., L. Novotna, A. Blount, and B.F. Lillemeier. 2017. A cycle of Zap70 kinase activation and release from the TCR amplifies and disperses antigenic stimuli. Nat Immunol. 18:86–95.

Keshvara, L.M., C.C. Isaacson, T.M. Yankee, R. Sarac, M.L. Harrison, and R.L. Geahlen. 1998. Syk- and Lyn-dependent phosphorylation of Syk on multiple tyrosines following B cell activation includes a site that negatively regulates signaling. J Immunol. 161:5276–5283.

Klammt, C., L. Novotna, D.T. Li, M. Wolf, A. Blount, K. Zhang, J.R. Fitchett, and B.F. Lillemeier. 2015. T cell receptor dwell times control the kinase activity of Zap70. Nat Immunol. 16:961–969.

Lin, J., M.J. Wester, M.S. Graus, K.A. Lidke, and A.K. Neumann. 2016. Nanoscopic cell-wall architecture of an immunogenic ligand in Candida albicans during antifungal drug treatment. Mol Biol Cell. 27:1002–1014.

Liu, Z.J., H. Haleem-Smith, H. Chen, and H. Metzger. 2001. Unexpected signals in a system subject to kinetic proofreading. Proc Natl Acad Sci U S A. 98:7289–7294.

Lombardo, L.J., F.Y. Lee, P. Chen, D. Norris, J.C. Barrish, K. Behnia, S. Castaneda, L.A. Cornelius, J. Das, A.M. Doweyko, C. Fairchild, J.T. Hunt, I. Inigo, K. Johnston, A. Kamath, D. Kan, H. Klei, P. Marathe, S. Pang, R. Peterson, S. Pitt, G.L. Schieven, R.J. Schmidt, J. Tokarski, M.L. Wen, J. Wityak, and R.M. Borzilleri. 2004. Discovery of N -(2-ch l oro-6-m ethyl- phenyl)-2-(6-(4-(2- hydroxyethyl)- piperazin-1-yl)-2-methylpyrimidin-4- ylamino)thiazole-5-carboxamide (BMS-354825), a dual Src/Abl kinase inhibitor with potent antitumor activity in preclinical assays. J Med Chem. 47:6658–6661.

Lopez, C.A., A. Sethi, B. Goldstein, B.S. Wilson, and GnanakaranS.. 2015. Membrane-mediated regulation of the intrinsically disordered CD3 cytoplasmic tail of the TCR. Biophys J. 108:2481–2491.

Lupher, M.L., Jr., N. Rao, N.L. Lill, C.E. Andoniou, S. Miyake, E.A. Clark, B. Druker, and H. Band. 1998. Cbl-mediated negative regulation of the Syk tyrosine kinase. A critical role for Cbl phosphotyrosine-binding domain binding to Syk phosphotyrosine 323. J Biol Chem. 273:35273–35281.

Mahajan, A., D. Barua, P. Cutler, D.S. Lidke, F.A. Espinoza, C. Pehlke, R. Grattan, Y. Kawakami, C.-S. Tung, A.R.M. Bradbury, W.S. Hlavacek, and B.S. Wilson. 2014. Optimal Aggregation of FcεRI with a Structurally Defined Trivalent Ligand Overrides Negative Regulation Driven by Phosphatases. ACS chemical biology.

Margolis, B., P. Hu, S. Katzav, W. Li, J.M. Oliver, A. Ullrich, A. Weiss, and J. Schlessinger. 1992. Tyrosine phosphorylation of vav proto-oncogene product containing SH2 domain and transcription factor motifs. Nature. 356:71–74.

McKeithan, T.W. 1995. Kinetic proofreading in T-cell receptor signal transduction. Proc Natl Acad Sci U S A. 92:5042–5046.

Menon, A.K., D. Holowka, W.W. Webb, and B. Baird. 1986. Cross-linking of receptor-bound IgE to aggregates larger than dimers leads to rapid immobilization. The Journal of cell biology. 102:541–550.

Metcalfe, D.D., D. Baram, and Y.A. Mekori. 1997. Mast cells. Physiol Rev. 77:1033–1079.

Metzger, H., G. Alcaraz, R. Hohman, J.P. Kinet, V. Pribluda, and R. Quarto. 1986. The receptor with high affinity for immunoglobulin E. Annu Rev Immunol. 4:419–470.

Murphy, K.P. 2012. Machine Learning: A Probabilistic Perspective. The MIT Press. 1096 pp.

O’Donoghue, G.P., R.M. Pielak, A.A. Smoligovets, J.J. Lin, and J.T. Groves. 2013. Direct single molecule measurement of TCR triggering by agonist pMHC in living primary T cells. eLife. 2:e00778.

Palacios, E.H., and A. Weiss. 2007. Distinct roles for Syk and ZAP-70 during early thymocyte development. J Exp Med. 204:1703–1715.

Park, M.J., R. Sheng, A. Silkov, D.J. Jung, Z.G. Wang, Y. Xin, H. Kim, P. Thiagarajan-Rosenkranz, S. Song, Y. Yoon, W. Nam, I. Kim, E. Kim, D.G. Lee, Y. Chen, I. Singaram, L. Wang, M.H. Jang, C.S. Hwang, B. Honig, S. Ryu, J. Lorieau, Y.M. Kim, and W. Cho. 2016. SH2 Domains Serve as Lipid-Binding Modules for pTyr-Signaling Proteins. Mol Cell. 62:7–20.

Pawitan, Y. 2001. In All Likelihood: Statistical Modelling and Inference Using Likelihood. OUP Oxford.

Pfeiffer, J.R., and J.M. Oliver. 1994. Tyrosine kinase-dependent assembly of actin plaques linking Fc epsilon R1 cross-linking to increased cell substrate adhesion in RBL-2H3 tumor mast cells. J Immunol. 152:270–279.

Presman, D.M., D.A. Ball, V. Paakinaho, J.B. Grimm, L.D. Lavis, T.S. Karpova, and G.L. Hager. 2017. Quantifying transcription factor binding dynamics at the single-molecule level in live cells. Methods.

Radhakrishnan, K., A. Halasz, M.M. McCabe, J.S. Edwards, and B.S. Wilson. 2012. Mathematical simulation of membrane protein clustering for efficient signal transduction. Ann Biomed Eng. 40:2307–2318.

Relich, P.K., M.J. Olah, P.J. Cutler, and K.A. Lidke. 2016. Estimation of the diffusion constant from intermittent trajectories with variable position uncertainties. Phys Rev E. 93:042401.

Sada, K., T. Takano, S. Yanagi, and H. Yamamura. 2001. Structure and function of Syk protein-tyrosine kinase. Journal of Biochemistry. 130:177–186.

Sanderson, M.P., E. Wex, T. Kono, K. Uto, and A. Schnapp. 2010. Syk and Lyn mediate distinct Syk phosphorylation events in FcεRI-signal transduction: implications for regulation of IgE-mediated degranulation. Molecular Immunology. 48:171–178.

Schnabel, R.B., and E. Eskow. 1999. A Revised Modified Cholesky Factorization Algorithm. SIAM Journal on Optimization. 9:1135–1148.

Schwartz, S.L., Q. Yan, C.A. Telmer, K.A. Lidke, M.P. Bruchez, and D.S. Lidke. 2015. Fluorogen-activating proteins provide tunable labeling densities for tracking FcepsilonRI independent of IgE. ACS Chem Biol. 10:539–546.

Shaner, N.C., G.G. Lambert, A. Chammas, Y. Ni, P.J. Cranfill, M.A. Baird, B.R. Sell, J.R. Allen, R.N. Day, M. Israelsson, M.W. Davidson, and J. Wang. 2013. A bright monomeric green fluorescent protein derived from Branchiostoma lanceolatum. Nat Methods. 10:407–409.

Shelby, S.A., D.A. Holowka, B.A. Baird, and S.L. Veatch. 2014. Super-Resolution Localization Microscopy Identifies Distinct Stages of Antigen-Induced IgE Receptor Cross-Linking and Immobilization in Rbl-2H3 Mast Cells. Biophysical Journal. 106:238a–238a.

Shiue, L., M.J. Zoller, and J.S. Brugge. 1995. Syk is activated by phosphotyrosine-containing peptides representing the tyrosine-based activation motifs of the high affinity receptor for IgE. J Biol Chem. 270:10498–10502.

Sigalov, A. 2005. Multi-chain immune recognition receptors: spatial organization and signal transduction. Seminars in Immunology. 17:51–64.

Simon, M., L. Vanes, R.L. Geahlen, and V.L.J. Tybulewicz. 2005. Distinct roles for the linker region tyrosines of Syk in FcepsilonRI signaling in primary mast cells. The Journal of biological chemistry. 280:4510–4517.

Smith, A.J., Z. Surviladze, E.A. Gaudet, J.M. Backer, C.A. Mitchell, and B.S. Wilson. 2001. p110beta and p110delta phosphatidylinositol 3-kinases up-regulate Fc(epsilon)RI-activated Ca2+ influx by enhancing inositol 1,4,5-trisphosphate production. J Biol Chem. 276:17213–17220.

Smith, C.S., N. Joseph, B. Rieger, and K.A. Lidke. 2010. Fast, single-molecule localization that achieves theoretically minimum uncertainty. Nature methods. 7:373–375.

Steinkamp, M.P., S.T. Low-Nam, S. Yang, K.A. Lidke, D.S. Lidke, and B.S. Wilson 2014 erbB3 is an active tyrosine kinase capable of homo- and heterointeractions. Mol Cell Biol. 34:965–977.

Suzuki, R., S. Leach, W. Liu, E. Ralston, J. Scheffel, W. Zhang, C.A. Lowell, and J. Rivera. 2014. Molecular editing of cellular responses by the high-affinity receptor for IgE. Science. 343:1021–1025.

Tierney, L., and J.B. Kadane. 1986. Accurate Approximations for Posterior Moments and Marginal Densities. J Am Stat Assoc. 81:82–86.

Torigoe, C., J.R. Faeder, J.M. Oliver, and B. Goldstein. 2007. Kinetic proofreading of ligand-FcepsilonRI interactions may persist beyond LAT phosphorylation. J Immunol. 178:3530–3535.

Tsang, E., A.M. Giannetti, D. Shaw, M. Dinh, J.K.Y. Tse, S. Gandhi, H. Ho, S. Wang, E. Papp, and J.M. Bradshaw. 2008. Molecular mechanism of the Syk activation switch. The Journal of biological chemistry. 283:32650–32659.

Turner, M., E. Schweighoffer, F. Colucci, J.P. Di Santo, and V.L. Tybulewicz. 2000. Tyrosine kinase SYK: Essential functions for immunoreceptor signalling. Immunology Today. 21:148–154.

Valley, C.C., D.J. Arndt-Jovin, N. Karedla, M.P. Steinkamp, A.I. Chizhik, W.S. Hlavacek, B.S. Wilson, K.A. Lidke, and D.S. Lidke. 2015. Enhanced dimerization drives ligand-independent activity of mutant epidermal growth factor receptor in lung cancer. Mol Biol Cell. 26:4087–4099.

van de Linde, S., A. Loschberger, T. Klein, M. Heidbreder, S. Wolter, M. Heilemann, and M. Sauer. 2011. Direct stochastic optical reconstruction microscopy with standard fluorescent probes. Nat Protoc. 6:991–1009.

van den Dries, K., S.L. Schwartz, J. Byars, M.B. Meddens, M. Bolomini-Vittori, D.S. Lidke, C.G. Figdor, K.A. Lidke, and A. Cambi. 2013. Dual-color superresolution microscopy reveals nanoscale organization of mechanosensory podosomes. Mol Biol Cell. 24:2112–2123.

Veatch, S.L., E.N. Chiang, P. Sengupta, D.A. Holowka, and B.A. Baird. 2012. Quantitative nanoscale analysis of IgE-FcepsilonRI clustering and coupling to early signaling proteins. J Phys Chem B. 116:6923–6935.

Vonakis, B.M., H. Haleem-Smith, P. Benjamin, and H. Metzger. 2001. Interaction between the unphosphorylated receptor with high affinity for IgE and Lyn kinase. J Biol Chem. 276:1041–1050.

Wilson, B.S., J.M. Oliver, and D.S. Lidke. 2011. Spatio-temporal signaling in mast cells. Adv Exp Med Biol. 716:91–106.

Wilson, B.S., J.R. Pfeiffer, and J.M. Oliver. 2000. Observing FcepsilonRI signaling from the inside of the mast cell membrane. The Journal of cell biology. 149:1131–1142.

Wilson, B.S., S.L. Steinberg, K. Liederman, J.R. Pfeiffer, Z. Surviladze, J. Zhang, L.E. Samelson, L.H. Yang, P.G. Kotula, and J.M. Oliver. 2004. Markers for detergent-resistant lipid rafts occupy distinct and dynamic domains in native membranes. Mol Biol Cell. 15:2580–2592.

Woody, M.S., J.H. Lewis, M.J. Greenberg, Y.E. Goldman, and E.M. Ostap. 2016. MEMLET: An Easy-to-Use Tool for Data Fitting and Model Comparison Using Maximum-Likelihood Estimation. Biophys J. 111:273–282.

Yamashita, T., S.Y. Mao, and H. Metzger. 1994. Aggregation of the high-affinity IgE receptor and enhanced activity of p53/56lyn protein-tyrosine kinase. Proc Natl Acad Sci U S A. 91:11251–11255.

Yan, Q., S.L. Schwartz, S. Maji, F. Huang, C. Szent-Gyorgyi, D.S. Lidke, K.A. Lidke, and M.P. Bruchez. 2014. Localization microscopy using noncovalent fluorogen activation by genetically encoded fluorogen-activating proteins. Chemphyschem. 15:687–695.

Yu, Y., S. Gaillard, J.M. Phillip, T.-C. Huang, S.M. Pinto, N.G. Tessarollo, Z. Zhang, A. Pandey, D. Wirtz, A. Ayhan, B. Davidson, T.-L. Wang, and I.-M. Shih. 2015. Inhibition of Spleen Tyrosine Kinase Potentiates Paclitaxel-Induced Cytotoxicity in Ovarian Cancer Cells by Stabilizing Microtubules. Cancer cell. 28:82–96.

Zhang, J., E.H. Berenstein, R.L. Evans, and R.P. Siraganian. 1996. Transfection of Syk protein tyrosine kinase reconstitutes high affinity IgE receptor-mediated degranulation in a Syk-negative variant of rat basophilic leukemia RBL-2H3 cells. J Exp Med. 184:71–79.

Zhang, Y., H. Oh, R.A. Burton, J.W. Burgner, R.L. Geahlen, and C.B. Post. 2008. Tyr130 phosphorylation triggers Syk release from antigen receptor by long-distance conformational uncoupling. Proceedings of the National Academy of Sciences of the United States of America. 105:11760–11765.

